# The oxidative stress response-related peroxiredoxin Tsa1b of *Candida auris* functions as a virulence factor that promotes infection

**DOI:** 10.1101/2024.11.26.625378

**Authors:** Maialen Areitio, Oier Rodriguez-Erenaga, Leire Aparicio-Fernandez, Lucía Abio-Dorronsoro, Leire Martin-Souto, Uxue Perez-Cuesta, Idoia Buldain, Beñat Zaldibar, Alba Ruiz-Gaitan, Javier Pemán, Salomé Leibundgut-Landmann, Aitor Rementeria, Aitziber Antoran, Andoni Ramirez-Garcia

## Abstract

The difficulty of accurately identifying *Candida auris* and the high resistance rates presented have increased the concern in the healthcare setting. Due to this, the aim of this study was to analyse the fungal response to oxidative stress. To achieve this goal, gene and protein expression were examined using qPCR and two-dimensional electrophoresis, respectively, peroxiredoxin Tsa1b being discovered to be overexpressed under oxidative stress. Besides, its antigenicity was also confirmed by western blotting. Subsequently, the significance of Tsa1b was next investigated by creating and characterizing the *C. auris ΔTSA1B* and *C. auris ΔTSA1B::TSA1B* strains using CRISPR-Cas9. The findings demonstrated that the *ΔTSA1B* strain was more susceptible to oxidative and cell wall stressors than the wild-type strain, which was consistent with an increase in the cell wall β-glucan amounts when grown in the presence of oxidative stress. Furthermore, the *ΔTSA1B* strain was also more vulnerable to the presence of dendritic cells and bone marrow-derived macrophages. Finally, *in vivo* infections performed in *Galleria mellonella* and mice showed a slower progression of the disease in those animals infected with the mutant strain. In conclusion, the peroxiredoxin Tsa1b has been identified as an important protein for the *C. auris* response to oxidative stress and as a virulence factor, allowing for a more thorough knowledge of the pathobiology of this yeast. This study points out the potential that this protein may have for the development of new diagnostic and therapeutic approaches.

**HIGHLIGHTS:**

- Several metabolic proteins are implicated in *C. auris* response to oxidative stress
- *C. auris* response to oxidative stress is influenced by the Tsa1b peroxiredoxin
- Lack of Tsa1b generates more susceptibility to stresses and an altered cell wall
- *C. auris* Tsa1b is involved in the fungal interaction with host immune cells
- The Tsa1b of *C. auris* contributes to the progression of the infection *in vivo*

## INTRODUCTION

The number of Invasive Fungal Infections (IFI) has been globally increasing in recent years. Nowadays, more than 300 million IFI cases/year are detected, with mortality rates ranging from 20% to almost 100% (Denning, 2024). The main infectious agents are *Candida* species (Magill et al., 2014), *Candida albicans, Candida glabrata* (recently renamed as *Nakaseomyces glabratus*), *Candida parapsilosis* and *Candida tropicalis* being the main causative ones (Kim et al., 2020; Webb et al., 2018). In addition, the infections caused by *Candida auris*, firstly identified in 2009 (Satoh et al., 2009), have gained importance over the last years (Du et al., 2020). As a result, this fungus has been the first fungal species for which the Centers for Disease Control and Prevention (CDC, USA) has issued a public health global alert (CDC, 2016), and it has been included on the Critical Group of the Fungal Pathogen Priority List published by the World Health Organization (WHO) (WHO, 2022).

One of the most significant virulence factors of *C. auris* is its resistance to stresses, which leds to inefficient decolonization of patients and abiotic surfaces (Spivak and Hanson, 2018), and worsens the prognosis of the patients (Schelenz et al., 2016). In fact, a higher growth capacity of *C. auris* than *C. albicans* has been observed in presence of oxidative compounds or disinfectants (Day et al., 2018; Heaney et al., 2020). Specifically, the resistance of *C. auris* to oxidative stress, mainly caused by Reactive Oxygen Species (ROS), decisively influences several processes related to pathogenicity, including resistance to decontamination processes and immune response (Cadnum et al., 2017; Herb and Schramm, 2021).

In order to reduce the effect of ROS, a rapid response is activated in the yeast by expressing antioxidant enzymes, including catalase (Cat1), glutathione peroxidase (Gpx), cytochrome C peroxidase (Ccp1), and superoxide dismutases (SODs), as well as components of the glutathione/glutaredoxin (Gsh1, Ttr1) and thioredoxin (Tsa1b, Trx1, Trr1) systems, which are important for the repair of proteins damaged by these molecules (Dantas et al., 2015). In addition, signaling proteins involved in the regulation of these enzymes, such as Hog1, Ssk1, and Mad2, might also be virulence factors of *C. auris*, as it happens in *C. albicans* (Bai et al., 2002; Chauhan et al., 2003; da Silva Dantas et al., 2010; Dantas et al., 2015).

Due to that, in this research the response of *C. auris* to oxidative stress generated by H_2_O_2_ was studied by gene and protein expression analyses using RT-qPCR and 2-Dimensional Electrophoresis (2-DE), respectively. The results showed that Tsa1b peroxiredoxin was the only molecule detected in both analyses. Thus, the implication of this protein on oxidative response and virulence was studied through the generation of a *C. auris* deletion mutant strain by CRISPR-Cas9 technique. The characterization of this mutant showed that Tsa1b is important not only for the resistance of *C. auris* to oxidative stress but also for the infectious process, which makes it a potential target for the development of novel therapeutic approaches.

## MATERIALS AND METHODS

### Microorganisms, cell cultures, and human sera

*Candida auris* clinical isolates CJ-194, CJ-195, CJ-196, CJ-197, and CJ-198, and DSM 105992 strain were used. The isolates CJ-197 and CJ-198 were deposited on the Spanish Type Culture Collection (CECT) as CECT 13225 and CECT 13226, respectively. In addition, *C. albicans* SC 5314 and CECT 13062, *Candida haemulonii* CECT 11935, and *C. parapsilosis* ATCC 22019 strains were also used. Besides, throughout the development of this study, *C. auris ΔTSA1B* and *C. auris ΔTSA1B::TSA1B* strains were constructed. *C. auris* isolates from bloodstream infections (CJ-194 - CJ-198) were obtained from the University and Polytechnic La Fe Hospital (Valencia, Spain). The Ethical Committee from the University of the Basque Country (UPV/EHU) (ref. M30/2020/019 and M30/2022/098) approved all procedures.

All the strains were cryopreserved at −80°C and cultured on Sabouraud Dextrose Agar (SDA) (Condalab, Madrid, Spain) at 37°C for 24 hours before use, and then, yeast cell densities were adjusted to inoculate 5 x 10^5^ yeasts/mL in Sabouraud Dextrose Broth (SDB) (Millipore, Massachusetts, USA), incubated at 37°C overnight at 120 rpm and collected by centrifugation (8,100 *g*, 3 minutes).

The dendritic cell (DC) line DC1940, which constitutively expresses green fluorescence protein (GFP) (Steiner et al., 2008), was kept in Iscove′s Modified Dulbecco′s Medium, supplemented with 10% decomplemented Fetal Calf Serum (FCS), 1% penicillin/streptomycin, 2 mM L-glutamine and 50 μM β-mercaptoethanol and incubated in humidified conditions at 37°C and 5% CO_2_.

Granulocyte-macrophage colony-stimulating factor (GM-CSF) needed for the differentiation of monocytes into macrophages was obtained from L929 cell line (ATCC CCL-1) culture supernatants. L929 cells were cultured in R10* media, consisting of Roswell Park Memorial Institute (RPMI) medium supplemented with 10% decomplemented FCS, 1% penicillin/streptomycin, 2 mM L-glutamine, 50 μM β-mercaptoethanol, 10 mM HEPES, 1% non-essential amino acids, and 1% sodium pyruvate, and incubated in humidified conditions at 37°C and 5% CO_2_ for 7 days. Then, the supernatant was filtered and stored at −20°C.

Monocyte cells obtained from the hind legs of 8-12 weeks old C57BL/6 mice were differentiated into bone-marrow derived macrophages (BMDMs) in R10* medium supplemented with 20% of L929 conditioned medium, as previously described (Clancey et al., 2020).

Sera from patients with disseminated *C. auris* infections were obtained from University and Polytechnic La Fe Hospital with approval of the Ethical Committee (ref. 2020-642-1).

### Characterization of *C. auris* clinical isolates

To analyse the growth of *C. auris* isolates on different environmental stresses, *Candida* spp. cells were collected as previously indicated. Then, cell densities were adjusted to 5 x 10^7^ cells/mL and 10 µl of 1:10 serial dilutions were plated on SDA or Yeast Peptone Dextrose (YPD) plates supplemented with 200 μg/mL of Calcofluor White (CFW), 300 μg/mL of Congo Red (CR), 0.05% of sodium dodecyl sulphate (SDS), 8 mM or 15 mM of H_2_O_2_, 1.75 M NaCl and 0.6 M potassium chloride (KCl). SDA or YPD plates with pH values of 3 or 10 were also prepared. The plates were incubated at 37°C for 24-48 hours.

The antifungal susceptibility testing (AFST) assay was performed as described in the European Committee on Antimicrobial Susceptibility Testing (EUCAST) E.Def 7.3.1 document. *C. parapsilosis* ATCC 22019 was used as the control strain.

### Gene expression analysis of *C. auris* in presence of oxidative stress

To obtain total RNA of *C. auris* alone or under conditions of oxidative stress, 5 x 10^5^ yeasts/mL were grown at 37°C and 120 rpm in SDB for 8, 16 or 24 hours. When required, SDB medium was supplemented with 8 mM or 15 mM of H_2_O_2_. Then, the culture was centrifuged at 11,419 *g* at 4°C for 10 minutes and washed with cold sterile distilled water. Total RNA was extracted following NZY Total RNA Isolation kit (NZYTech, Lisbon, Portugal) guidelines, with a few modifications. Briefly, yeast cells were lysed with 0.1 µm diameter zirconium beads, using a Millmix 20 Bead-Beater (Thetnica, Železniki, Slovenia) and three cycles of lysis at 30 Hz for 1 minute with a posterior incubation on ice for 1 minute. RNA integrity and concentrations were verified by electrophoresis on agarose gels supplemented with 1% bleach and using the NanoDropTM Lite (Thermo Fisher Scientific, Waltham, USA) spectrophotometer, respectively.

Gene expression analysis of 10 *C. auris* genes (*CAT1, SOD1, SOD2, SOD6, HOG1, MAD2, TSA1B, HYR1, SSK1* and *CCP1*) homologous to those previously described as relevant on the oxidative stress response carried out by *C. albicans* (Dantas et al., 2015) was performed for *C. auris* CECT 13225 and DSM 105992 in absence and presence of 8 mM or 15 mM of H_2_O_2_. RNA samples obtained from the fungal cells were converted to cDNA with NZY First-Strand cDNA Synthesis Kit (NZYTech) and analysed by the method previously described (Aparicio-Fernandez et al., 2024). The primers used are listed in **Table S1**. Fold change (FC) values relative to the *C. auris ACT1* housekeeping gene were calculated by the Livak method (2^-ΔΔCq^).

### Total protein extract obtention and separation by electrophoresis

Total protein extract was obtained as previously described (Areitio et al., 2024), with supplementation of SDB medium with 8 mM or 15 mM of H_2_O_2_ when required. Protein concentration was measured using Pierce 660 nm Protein Assay Reagent (Thermo Fisher Scientific, Waltham, MA, USA).

Protein separation was made by one-dimensional electrophoresis (1-DE) or 2-DE. The 1-DE was carried out in 10% acrylamide gels and running them at 70 mA, 100 W, and 200 V for 45 min in a Miniprotean II (Bio-Rad, Hercules, California, USA). For 2-DE, the method previously described (Pellon et al., 2016) was used for protein precipitation and separation. Page Ruler Plus (Thermo Fisher Scientific) was used as the protein standard. The gels were stained with Coomassie Brilliant Blue (CBB) and digitalized with ImageScanner III (GE Healthcare, Chicago, IL, USA) software or transferred to membranes for antigenic detection. All the 1-DE and 2-DE were made in triplicate, and only the most informative gels are shown.

The determination of overexpressed points was made based on the volume percentage (Vol%) parameter given by ImageMaster 2D Platinum Software (GE Healthcare). The spots detected only in all the replicates obtained from the oxidative stress condition and those presenting a mean Vol% at least four times higher than in the control condition were selected as overexpressed and subjected to identification.

### Identification of proteins

The selected protein spots were manually extracted from CBB stained gels and identified by liquid chromatography with tandem mass spectrometry (LC-MS/MS) in the SGIker proteomic services of the UPV/EHU, as previously described (Buldain et al., 2016), with a few modifications. LC-MS/MS was carried out on Exploris 240 mass spectrometer (Thermo Fisher Scientific) interfaced with EASY nLC-1200 system (Thermo Fisher Scientific). The search and identification of proteins was done as previously described (Areitio et al., 2024). The mass spectrometry proteomics data have been deposited to the ProteomeXchange Consortium via the PRIDE partner repository (Perez-Riverol et al., 2022) with the data set identifier PXD051894.

### Bioinformatic analysis of the identified proteins

The softwares DeepLoc 2.0 (https://services.healthtech.dtu.dk/services/DeepLoc-2.0/), SignalP 6.0 (https://services.healthtech.dtu.dk/services/SignalP-6.0/), SecretomeP 2.0 (https://services.healthtech.dtu.dk/services/SecretomeP-2.0/) and FaaPred (https://bioinfo.icgeb.res.in/faap/query.html) were used to predict the location, the secretion through signal peptides or non-classical methods, and the possible adhesion properties of the antigens, respectively. On DeepLoc 2.0 the cellular location was indicated. On the rest of the prediction pages, results were considered positive when the score was > 0.5 for SignalP 6.0, > 0.6 for SecretomeP 2.0, or setting a −0.8 threshold value for FaaPred. For the study of the functionality, family, and domains, the Interpro (https://www.ebi.ac.uk/interpro/) database was used. For the antigenicity and the protective antigen predictions, AntigenPRO (https://scratch.proteomics.ics.uci.edu/) and VaxiJen 2.0 (http://www.ddg-pharmfac.net/vaxijen/VaxiJen/VaxiJen.html) were used. AllerTOP v.2 (http://www.ddg-pharmfac.net/AllerTOP/) and AllergenFP (http://www.ddg-pharmfac.net/AllergenFP/) were chosen for the allergenicity prediction. Finally, NCBI BlastP (https://blast.ncbi.nlm.nih.gov/Blast.cgi) was used for the study of the homology and the alignment of the proteins with other fungal species, as well as with humans. Protein interaction networks studies were performed using the STRING tool (v12; https://string-db.org/).

### Protein isolation and purification

After analysing the proteome of the fungus under H_2_O_2_ presence, the selected protein spots were excised from the 2-DE gels by electroelution. Multiple replicates of the same spot were used until the appropriate concentration was achieved. Proteins were electroeluted with electroelution buffer (25 mM Tris, 192 mM glycine, 3.5 mM SDS, pH 8.3) using a 422 Electroeluter (Bio-Rad) system at room temperature (RT), at 60 mA for 5 hours, and stored at - 20°C.

### Antigen detection

For antigen detection, protein spots were transferred to Amersham Hybond-P poly(vinylidene difluoride) (PVDF) membranes (GE Healthcare) and the Western Blot (WB) technique was performed, as previously described (Areitio et al., 2024). Briefly, human serum diluted to 1/200 was used as primary antibody and human anti-IgG-HRP diluted to 1/100,000 was added as secondary antibody. The detection of immunoreactive proteins was achieved by ECL Advanced (NZYTech) in the G:BOX Chemi System (Syngene, Cambridge, UK) and ImageMaster 2D Platinum Software (GE Healthcare) was used for WB analysis.

When needed, the oxidation of glycoproteins transferred to the PVDF membrane was achieved by 50 mM sodium metaperiodate treatment in 100 mM sodium acetate buffer (pH 5.5) at RT for 30 minutes, and after washing the membrane, the WB process described above was followed.

### Generation of the knock-out and complemented *C. auris* strains by CRISPR-Cas9 method

The DNA from the *C. auris* CECT 13225 and DSM 105992 were obtained following a previously described protocol for *C. albicans* (Lõoke et al., 2017). For bacterial plasmid DNA extraction, which contained the hygromycin cassette, overnight liquid cultures of *Escherichia coli* were lysed and treated following GeneJET Plasmid Miniprep kit (Thermo Fisher Scientific) guidelines.

The crRNAs were designed in the adjacent 20 nucleotides of the desired protospacer adjacent motifs (PAM) recognized by the Cas9 enzyme **(Table S1)**, and obtained from Integrated DNA Technologies (Iowa, USA). The deletion mutant (*ΔTSA1B*) and complemented (*ΔTSA1B::TSA1B*) strains of the *TSA1B* gene were generated following a protocol previously described (Rybak et al., 2019). For *C. auris ΔTSA1B* strain generation, primer pair 1-2 were used **(Table S1)**, and to obtain the complemented strain primer pair 3-4 and 5-6 were used **(Table S1) (Figure 1)**. The NEBNext High-Fidelity polymerase (New England Biolabs, Ipswich, MA, USA) was used for the amplification of the cassettes.

**Figure 1.**
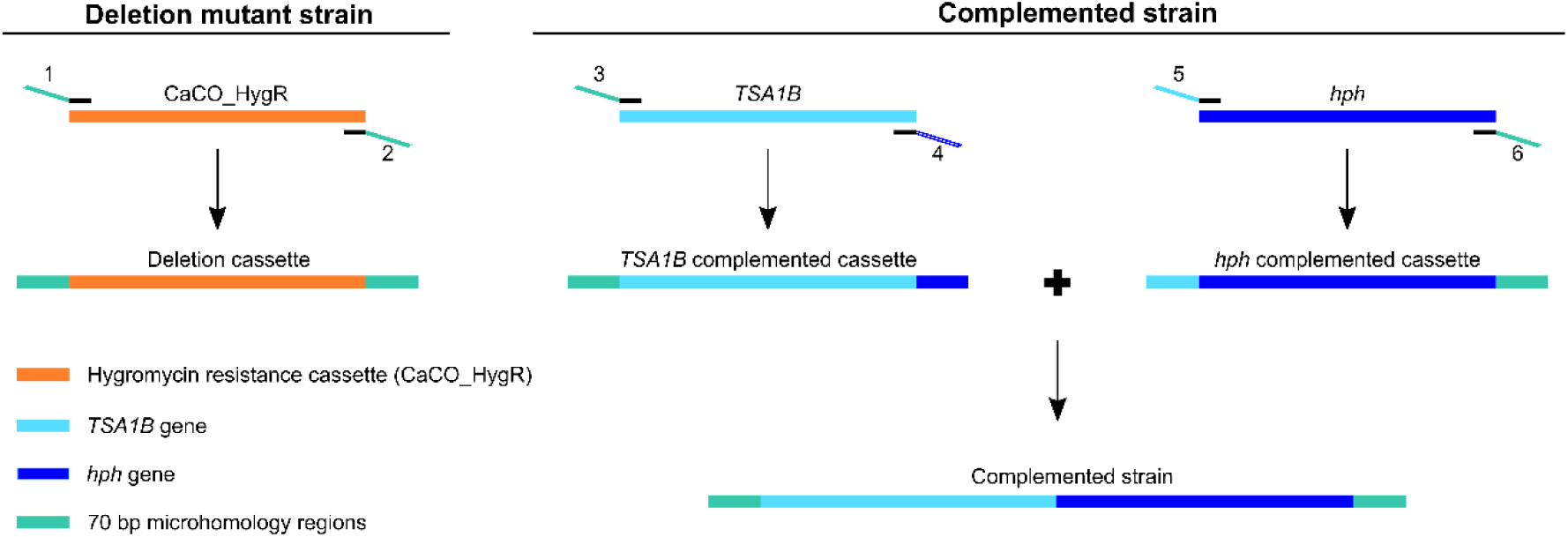
Scheme of the construction of the cassettes generated for the transformation of *Candida auris* strains. The CaCO_HygR cassette used for the construction of the deletion strain and the two cassettes used for the construction of the complemented strain are represented.

Only colonies of yeast cells that grew on YPD plates supplemented with hygromycin were tested and potential deletion mutant colonies were grown in Yeast Peptone Maltose (YPM) plates to induce the excision of the deletion cassette, as previously described (Rybak et al., 2019). The DreamTaq DNA polymerase (Thermo Fisher Scientific) with primer pairs a-b and c-d **(Table S1)** was used and results were confirmed by Sanger sequencing (Eurofins, Nantes, France).

### *C. auris* chitin and β-glucan quantification

Cell wall components chitin and β-glucan of *C. auris* were quantified by flow cytometry as previously described (Lee et al., 2018), with a few modifications. Briefly, after growing the fungus in absence and presence of 15 mM of H_2_O_2_, chitin and β-glucan staining was performed with 6 µg/mL of CFW (MP Biomedicals, Santa Ana, CA, USA) and 100 mg/mL of Aniline Blue (Sigma-Aldrich), respectively. Excitation was set at 405 nm for both compounds, and emission at 450 nm and 525 nm for CFW and aniline blue staining, respectively. A total of 10,000 cells were passed by CytoFLEX SRT (Beckman Coulter, Brea, CA, USA) and the obtained Mean Fluorescence Intensity (MFI) was analysed by CytExpert (Beckman Coulter) software.

### Interaction of *C. auris* with immune cells

Co-incubation of fungal and mammalian cells was done in R10* medium with a multiplicity of infection (MOI; *C. auris*:immune cells) of 5, for 30 minutes, 4 or 24 hours at 37°C and 5% CO_2_. After 4 hours of co-incubation, the survival of *C. auris* upon contact with immune cells and the ROS amount measurement of immune cells were determined. For the fungal cell survival assay, NP-40 detergent solution at 0,05% was added to lyse the immune cells and lysates were plated on YPD and Colony Forming Units (CFUs) were counted after incubation at 37°C for 24-48 hours. For ROS quantification, supernatants were removed and 5 µM dihydroethidium were added for 20 minutes at 37°C. After removal of supernatants, PBS was added, and fluorescence was immediately read at the Infinite 200 plate reader (Tecan, Männedorf, Switzerland) with excitation at 500 nm and emission at 580 nm. MFI values obtained after subtracting the blank were indicated.

After co-incubation for 4 and 24 hours, the survival of host immune cells upon contact with *C. auris* was measured. To achieve that, supernatants were collected, and Cytotoxicity detection Kit (LDH) (Sigma-Aldrich) was used to measure lactate dehydrogenase (LDH) release. Absorbance was measured at the Infinite 200 plate reader at 492 nm and 630 nm (control value). Cytotoxicity was calculated using the following equation: 100 x (infected cells – uninfected cells)/(lysed cells – uninfected cells). Uninfected cells and cells treated with 1% Triton-X were considered as negative and positive controls for lysis of host immune cells, respectively.

To determine the uptake capacity of *C. auris* by host immune cells, co-incubations of 30 minutes were performed, at two different conditions. For each experiment, two 96-well plates were set up, one at 37°C and 5% CO_2_, to allow attachment and internalization of fungal cells, and the other one at 4°C, to allow just attachment of *C. auris* to the immune cells surface. After co-incubation for 30 min, cells were detached and washed with fluorescence-activated cell sorting (FACS) buffer (PBS supplemented with 1% FCS, 5 mM EDTA, and 0.02% NaN_3_) three times by centrifugation at 2,855 *g* for 5 minutes. Then, surface antibodies (BioLegend, San Diego, CA, USA) specific for DC or macrophage antigenic markers were used for immune cell staining **(Table S2)**, at 4°C for 40 minutes. Samples were washed with FACS buffer and mammalian cells were permeabilized for 20 minutes at 4°C with BD Fix/Perm solution (BD Biosciences, Franklin Lakes, NJ, USA) and washed with 1X BD Perm/Wash buffer (BD Biosciences). *C. auris* was stained by addition of CFW 1:1000 dilution in PBS and incubated for 20 minutes at RT. Finally, samples were washed twice with PBS and resuspended in FACS buffer. Data was acquired on a SP6800 Spectral Analyzer (Sony, Minato, Tokyo, Japan) and analyzed using FlowJo software (BD Biosciences). Fungal uptake by immune cells was determined after gated data of fungal cells incubated at 4°C condition was subtracted from the gated data at 37°C condition. To gate DCs, events double positive for CD11c and GFP markers were selected, and for BMDMs, positive events for CD64. Gating of fungal cells was performed by selection of positive events for CFW staining.

Finally, gene expression analysis of *C. auris* genes previously identified in the gene expression analysis in presence of oxidative stress was performed after 2 and 4 hours of co-incubation. To obtain total RNA of *C. auris* cells, first, fungal cells were disrupted with 0.5 mm glass beads (Sigma-Aldrich, St Louis, MO, USA) and Tissue Lyzer (Qiagen) at 30 Hz for 2 minutes and, afterwards, 30 seconds of incubation on ice, repeating the process twice. After fungal lysis, RNeasy Mini Kit (Qiagen, Hilden, Germany) was used and RNA samples were converted to cDNA with RevertAid H Minus Reverse Transcriptase kit (Thermo Fisher Scientific) and analysed as previously described (Lemberg et al., 2022). The primers used are listed in **Table S1**. Expression levels were calculated relative to the *C. auris ACT1* housekeeping gene.

### Galleria mellonella infection

Sixth-instar larvae (Reptimercado, Murcia, Spain) were injected with 2.5 x 10^6^ cells of *C. auris* wild-type (WT), *ΔTSA1B* or *ΔTSA1B::TSA1B* strains in 10 µL PBS through the last right pro-leg. Sixteen larvae were used per group, and non-injected and PBS injected control groups were included. Larvae were incubated at 37°C and checked daily over 7 days. Three independent experiments were performed (n ≥ 48).

### Murine model of disseminated infection

Six-to eight-week-old Swiss female and male mice bred and maintained at the SGIker Animal Facility of the UPV/EHU were used. Animals were maintained with water and food *ad libitum* in filter-aerated sterile cages. The Animal Experimentation Ethical Committee from the UPV/EHU approved all the procedures (ref. M20/2022/099).

Mice were anesthetized by intraperitoneal injection of 100 mg/kg of ketamine and 10 mg/kg of xylazine. All infections were made by intravenous injection of fungal cells suspended in 0.2 mL of DPBS in the tail vein of each animal.

Forty mice were separated in four groups consisting of 10 mice each (five males and five females), on which Dulbecco’s Phosphate-Buffered Saline (DPBS), *C. auris* WT, *ΔTSA1B* and *ΔTSA1B::TSA1B* strains were injected. Control mice received 0.2 mL of DPBS intravenously, while infected mice were intravenously inoculated with 5 x 10^7^ yeasts/animal. Twenty-eight days after the injection, mice were euthanized and total blood as well as brain, lungs, kidneys, spleen, and liver were extracted. Organs were divided into two halves; one was used for fungal load determination through CFU counting, and the other one for the histological study, as previously described (Areitio et al., 2024), with few modifications. For general observation of tissues, Giemsa staining was used, as described by Martoja and Martoja-Pierson (1970).

### Statistics

Statistical analysis was performed using SPSS statistical software (version 28.0.1.1; Professional Statistic, Chicago, IL, USA). All the experiments were performed at least three times. Normality of data was checked using the Shapiro-Wilks test and homogeneity of the variance using the Levene test. Outlier detection was made by GraphPad Outlier Calculator tool (https://www.graphpad.com/quickcalcs/Grubbs1.cfm). Data were analysed by ANOVA method followed by multiple comparisons corrected with Dunnett’s test, unpaired t-test, and log-rank Mantel-Cox test. All the graphics were plotted using GraphPad Prism version 8 (GraphPad Prism Software Inc., San Diego, CA, USA). Values of p < 0.05 were considered statistically significant.

## RESULTS

### Characterization of *C. auris* isolates

The growth of the five *C. auris* clinical isolates CJ-194, CJ-195, CJ-196, CECT13225, and CECT 13226 was compared, in the presence of stressor compounds, to two *C. albicans* isolates (SC 5314 and CECT 13062), the most commonly isolated species in candidemia, and *C. haemulonii* CECT 11935 isolate, the phylogenetically most closely related species. Overall, similar or greater growth of *C. auris* than *C. albicans* and *C. haemulonii* was observed in most of the tested conditions **(Figure 2)**. Specifically, all *C. auris* isolates showed greater resistance to H_2_O_2_ than *C. albicans* and *C. haemulonii*, and the same trend was observed for SDS. Moreover, some *C. auris* isolates grew better than *C. albicans* and *C. haemulonii* in the presence of CFW and CR. On the rest of the four conditions tested, almost no differences were observed among the species, except for pH 3, where less growth of *C. auris* than *C. albicans* was observed.

**Figure 2.**
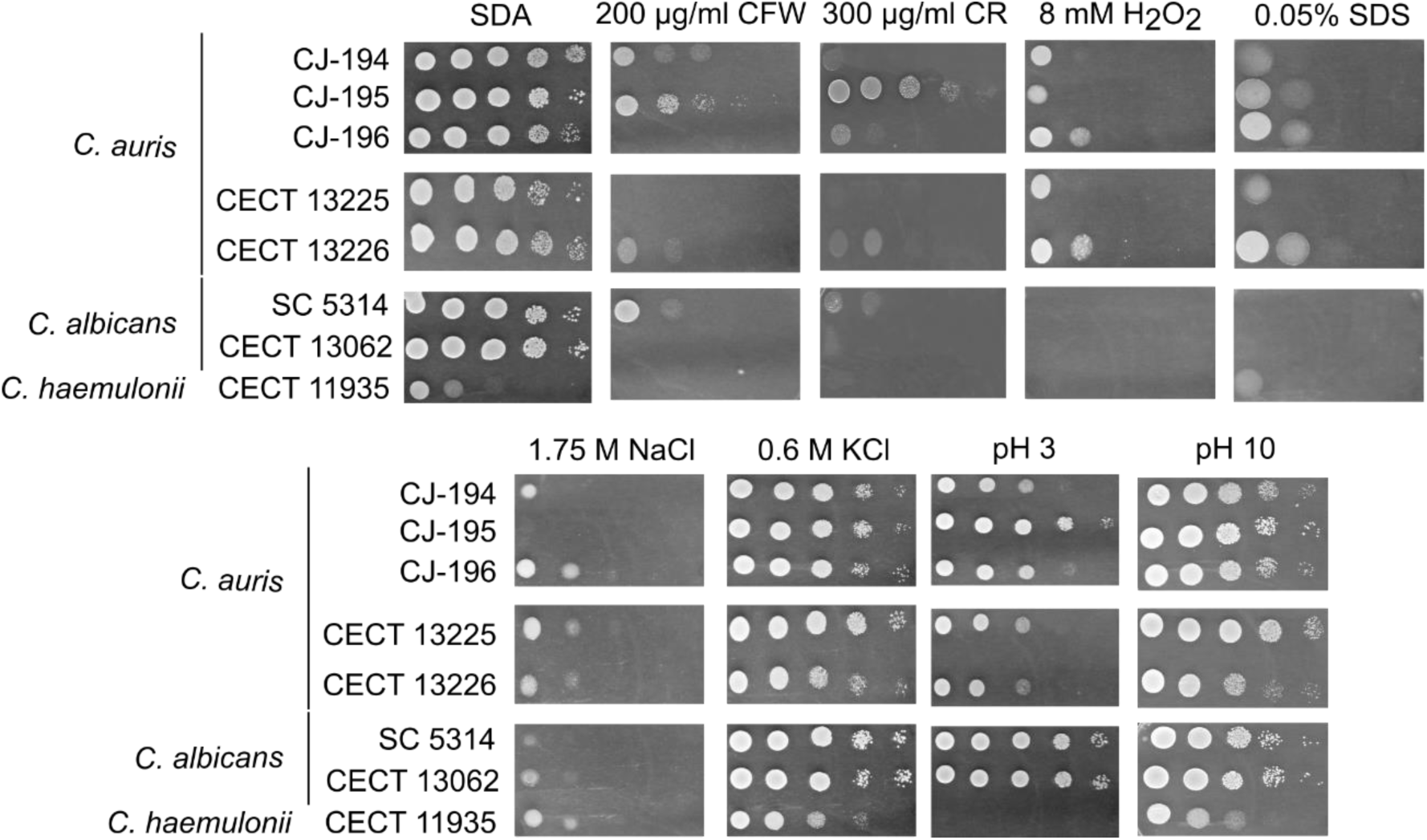
Growth of *Candida auris, Candida albicans* and *Candida haemulonii* strains in the presence of stressor compounds. Ten µl of 1:10 serial dilutions starting from 5 x 10^7^ yeasts/mL were plated on Sabouraud Dextrose Agar plates supplemented with 200 µg/mL CFW, 300 µg/mL CR, 0.05% SDS, 8 mM H2O2, 1.75 M NaCl, 0.6 M KCl and with pH 3 or pH 10 and incubated at 37°C for 24-48 hours. The most representative replicate of each condition is shown.

Besides, Minimal Inhibitory Concentration (MIC) values were determined using the EUCAST protocol and, according to the MIC values established by the CDC, the five *C. auris* isolates were resistant to fluconazole but not to amphotericin B or micafungin **(Table S3)**. In view that the isolate CECT 13225 showed the highest MIC values against amphotericin B and micafungin, it was selected to perform the rest of the experiments.

### Analysis of the expression of genes associated with the oxidative stress response in *C. auris*

The expression of the *C. auris* genes *CAT1, SOD1, SOD2, SOD6, HOG1, MAD2, TSA1B, HYR1, SSK1* and *CCP1* was studied by RT-qPCR. As shown in **Figure 3**, five out of the ten genes studied, *CAT1*, *SOD1*, *SOD6*, *TSA1B*, and *CCP1*, were overexpressed at 8 hours of incubation, four of them, *CAT1*, *SOD1*, *TSA1B* and *CCP1*, presenting values of fold change greater than 1.5 also at 16 hours. On the other hand, at 24 hours, underexpression of *CAT1, SOD2, SOD6, HYR1* and *SSK1* genes was observed, among which two of the genes previously overexpressed (*CAT1* and *SOD6*) were present.

**Figure 3.**
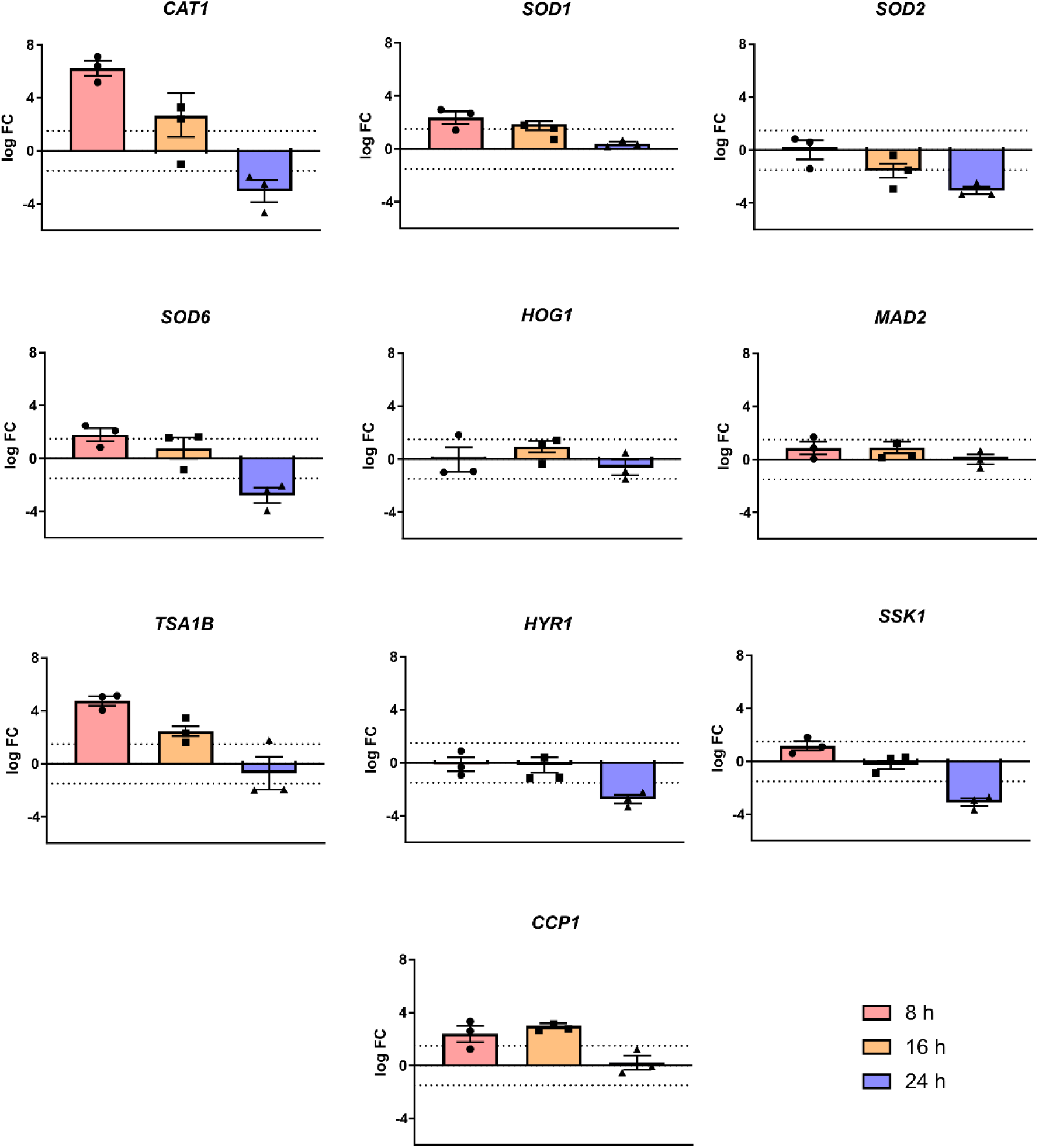
Expression analysis of genes related with the oxidative stress response on *Candida auris* after incubation with 8 mM H2O2 for 8, 16 and 24 hours. Analysis of *CAT1*, *SOD1*, *SOD2*, *SOD6*, *HOG1*, *MAD2*, *TSA1B*, *HYR1*, *SSK1* and *CCP1* genes of *C. auris* CECT 13225 was performed by RT-qPCR, by representing the fold change (FC) with respect to the control conditions and taking the reference expression of the constitutively expressed gene *ACT1*. Thresholds of 1.5 and - 1.5 were chosen for the determination of the overexpression or underexpression of the genes, respectively. Three biological replicates are represented for each condition. Mean and standard error of mean values are indicated.

### Identification and characterization of the *C. auris* overexpressed proteins under oxidative conditions

The proteomic changes caused by oxidative stress generated by H_2_O_2_ in *C. auris* were analysed by 2-DE. About 350 spots were studied in each condition, being differentially located depending on the sample, in the range of 25 - 100 kDa Mr and p*I* 5 - 8 in the control samples, and 25 - 130 kDa Mr and p*I* 5 - 8 in the H_2_O_2_-treated samples. Then, the 13 most overexpressed spots of the oxidative stress condition were selected to be identified by LC-MS/MS **(Figure 4)**.

**Figure 4.**
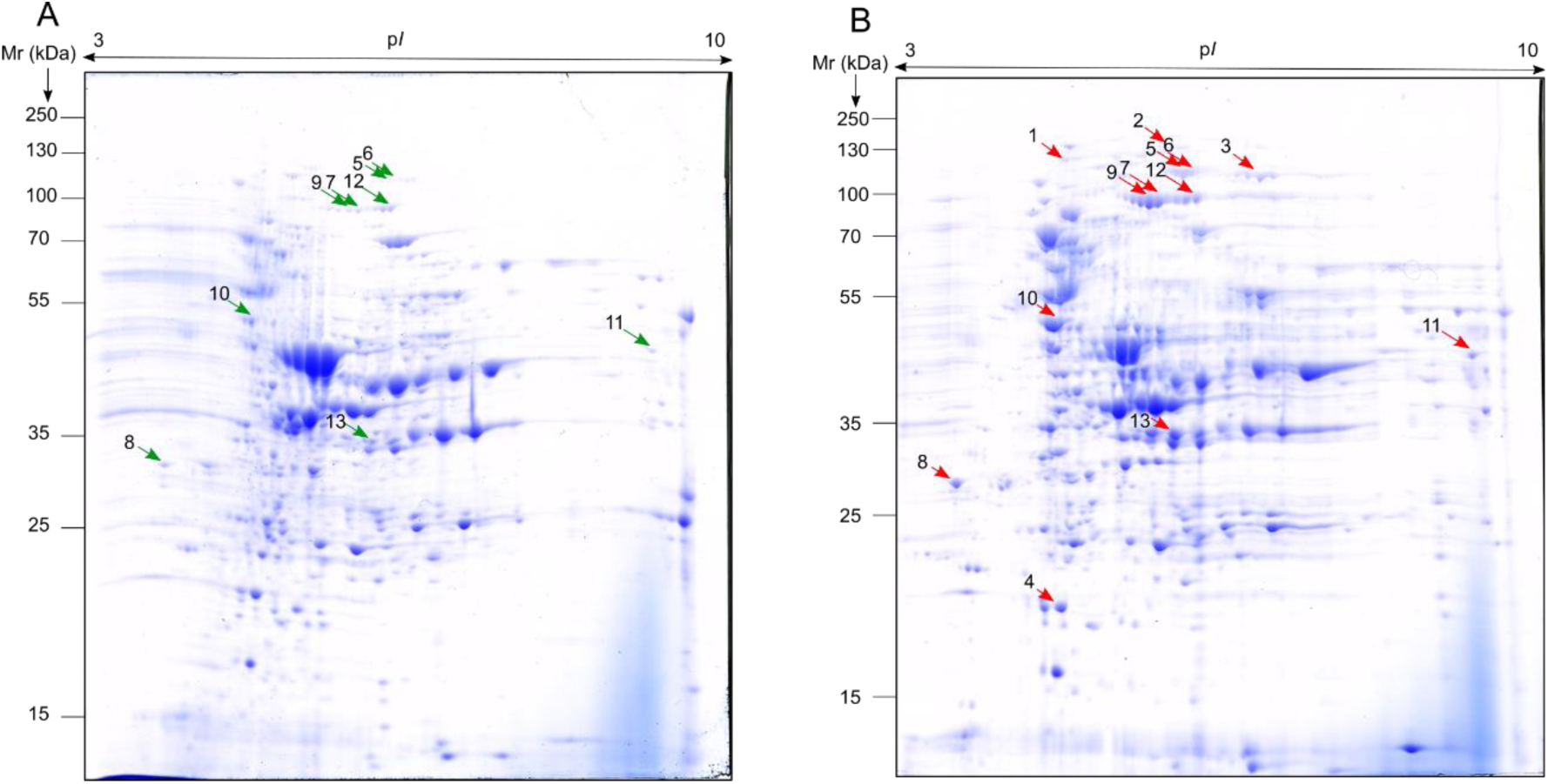
2-DE comparative analysis of *Candida auris* proteomes obtained in absence (control) and presence of oxidative stress. Proteomes of *C. auris* CECT 13225 obtained in the absence (A) and presence (B) of 8 mM of H2O2. Overexpressed points in H2O2 presence are indicated by red arrows; in green, points corresponding to red points in the proteome without oxidative stress.

The identified spots **(Table 1)** corresponded to the proteins ubiquitin-activating enzyme e1 (Uba1), elongation factor 3 (Cef3), leucine—tRNA ligase (Cdc60), thioredoxin-dependent peroxiredoxin (Tsa1b), mitochondrial C-1-tetrahydrofolate (Mis1), 5-methyltetrahydropteroyltriglutamate--homocysteine S-methyltransferase (Met6), 40S ribosomal protein S6 (Rps6), 60S ribosomal protein L8 (Rpl8), 40S ribosomal protein S4 (Rps4), elongation factor 1-α (Tef1), ATP synthase subunit β (Atp2), acetyl-CoA C-acetyltransferase (Erg10), mitochondrial aconitate hydratase (Aco1) and malate dehydrogenase (Mdh1). In addition to them, an uncharacterized protein was detected on spots 3 and 6, which had the same amino acid sequence as the formyltetrahydrofolate synthetase (Fau1). This name and abbreviation will, therefore, be used for it from now on. In addition, Met6 from spots 7, 9 and 12 revealed putative protein isoforms with the same Mr but different p*I*.

**Table 1.** Identification by LC-MS/MS of the overexpressed proteins of *Candida auris* when grown on conditions of oxidative stress. Information about Uniprot number, name of the protein, unique peptides identified, identified coverage, score of the identification, theoretical and experimental p*I* and Mr (kDa) values and VolH2O2%/VolCtrl% relationship of the identified spots are provided.

| Spot number | Uniprot number | Name of the protein | Unique Peptides | Cover (%) | Score | Theoretical pI/Mr (kDa) | Experimental pI/Mr (kDa) | Vol<sub>H2O2</sub>%/Vol<sub>Ctrl</sub>% |
| --- | --- | --- | --- | --- | --- | --- | --- | --- |
| 1 | A0A0L0NTT2 | Ubiquitin-activating enzyme e1 | 23 | 23 | 78.79 | 5.2/113.3 | 4.92/132.5 | - |
|  | A0A2H0ZKV5 | Elongation factor 3 | 18 | 18 | 68.92 | 6.04/116.4 |  |  |
| 2 | A0A2H0ZYD5 | Leucine--tRNA ligase | 15 | 12 | 37.95 | 6.1/124.3 | 6.11/152.3 | - |
| 3 | A0A2H0ZCS4 | Uncharacterized protein | 20 | 19 | 85.14 | 6.2/101.6 | 7.14/111.9 | - |
| 4 | A0A5Q7YJ27 | Thioredoxin-dependent peroxiredoxin | 6 | 31 | 56.83 | 5.16/21.7 | 4.73/21.1 | - |
| 5 | A0A0L0P7E6 | C-1-tetrahydrofolate mitochondrial | 13 | 13 | 39.66 | 6.2/101.7 | 6.22/117.8 | 10.33 |
| 6 | A0A2H0ZCS4 | Uncharacterized protein | 13 | 15 | 59.05 | 6.2/101.6 | 6.12/118.5 | 10.28 |
| 7 | A0A2H1A7R9 | 5-methyltetrahydropteroyltriglutamate--homocysteine S-methyltransferase | 37 | 34 | 682.3 | 5.91/86.1 | 5.92/93.2 | 10.09 |
| 8 | A0A2H1A5E3 | 40S ribosomal protein S6 | 6 | 31 | 25.32 | 10.29/27 | 3.68/31.1 | 7.23 |
|  | A0A890D5F9 | 60S ribosomal protein L8 | 7 | 26 | 20.76 | 10.15/29.1 |  |  |
|  | A0A510NYJ1 | 40S ribosomal protein S4 | 7 | 23 | 18.06 | 10.1/29.6 |  |  |
|  | A0A2H0ZIS9 | Elongation factor 1-alpha | 6 | 12 | 15.95 | 8.95/50 |  |  |
| 9 | A0A2H1A7R9 | 5-methyltetrahydropteroyltriglutamate--homocysteine S-methyltransferase | 22 | 24 | 102.99 | 5.91/86.1 | 5.8/92.2 | 6.68 |
| 10 | A0A890CXZ5 | ATP synthase subunit beta | 17 | 41 | 294.79 | 5.43/54.1 | 4.64/52.7 | 5.23 |
| 11 | A0A0L0P6W1 | Acetyl-CoA C-acetyltransferase | 14 | 37 | 93.65 | 8.02/41.4 | 9.12/47.7 | 4.73 |
| 12 | A0A510P142 | Aconitate hydratase, mitochondrial | 13 | 16 | 55.85 | 6.58/85 | 6.17/96.1 | 4.60 |
|  | A0A2H1A7R9 | 5-methyltetrahydropteroyltriglutamate--homocysteine S-methyltransferase | 15 | 19 | 46.26 | 5.91/86.1 |  |  |
| 13 | A0A5C1DVH2 | Malate dehydrogenase | 12 | 59 | 265.05 | 6.81/34.5 | 6.11/35 | 4.07 |

The bioinformatic analysis showed that 73% of the proteins identified were predicted to be located in the cytoplasm (Uba1, Cef3, Cdc60, Fau1, Tsa1b, Mis1, Met6, Rps6, Rpl8, Rps4, Tef1), whereas only a few of them in the mitochondria (Atp2, Aco1 and Mdh1), nucleus (Cdc60 and Met6), or peroxisome (Erg10). Furthermore, 20% of the proteins (Mdh1, Erg10 and Tef1) had adhesin function, and only the Rps4 was predicted to be secreted **(Figure 5A)**. In addition, 87% showed antigenic capacity (all except Rps4 and Atp2) and were predicted to be protective antigens (all but Cdc60 and Tsa1b). However, only Met6 seemed to possess allergenicity capacity **(Figure 5B)**.

**Figure 5.**
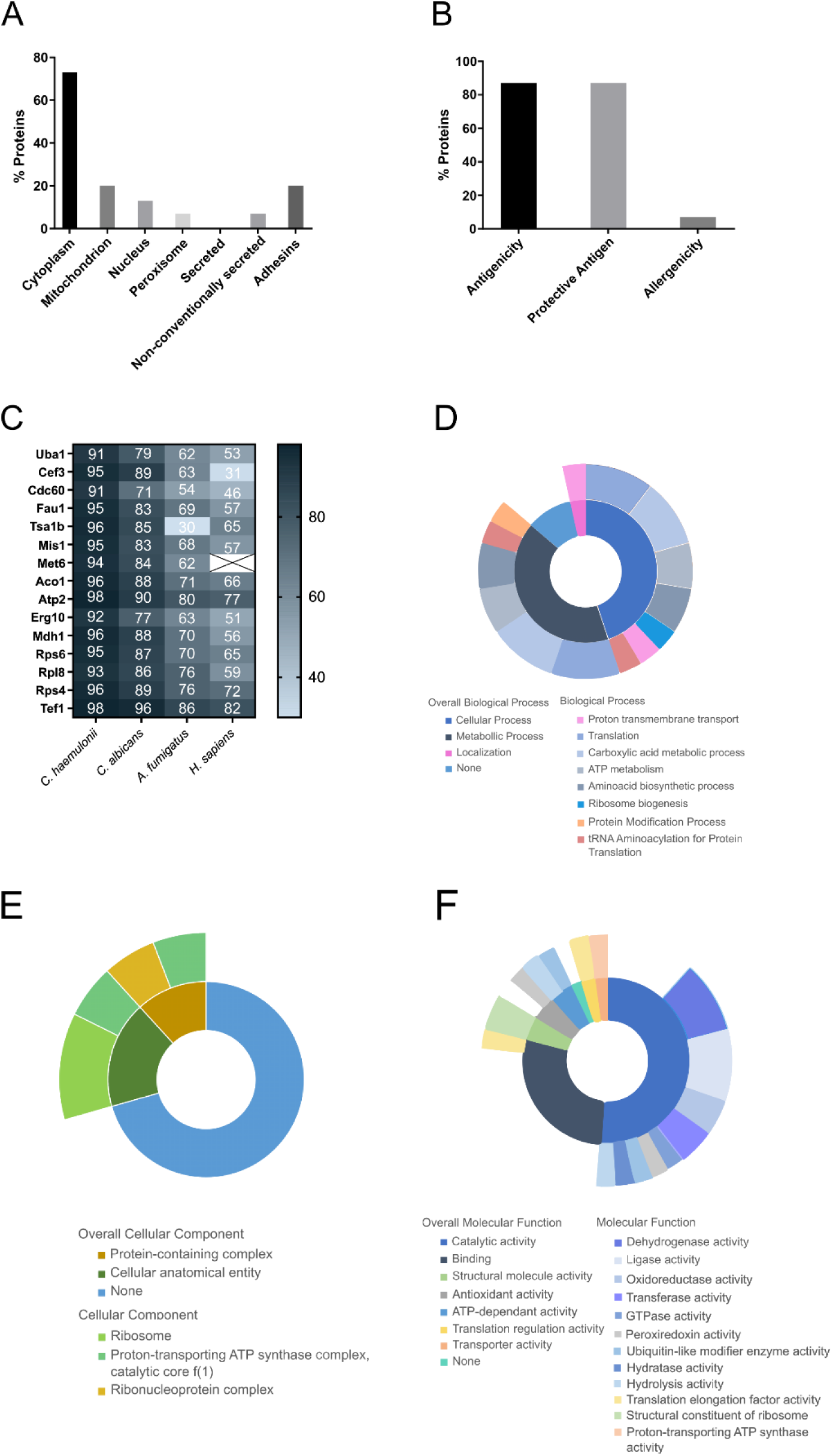
Bioinformatic analysis and study of the functionality of overexpressed *Candida auris* proteins on oxidative stress condition. Cellular localization and secretion (A), antigenicity, protective antigen capacity and allergenicity (B), homology values with *Candida haemulonii*, *Candida albicans*, *Aspergillus fumigatus* and *Homo sapiens*, and involvement of the identified proteins on biological processes (D), molecular functions (E) and cellular components (F) are indicated. The bioinformatic tools used for the analysis of the overexpressed proteins are listed on the Material and methods section.

As expected, given the phylogenetic relationship between the species, high homology was observed between the sequences of the proteins of *C. auris* and *C. haemulonii* (range 91% - 98%). Similarly, high percentages of homology were also observed with *C. albicans* (range 71% - 96%). On the other hand, lower homology percentages were found with *Aspergillus fumigatus,* and the lowest with *Homo sapiens* (range 30% - 86%) **(Figure 5C)**.

The study of the functionality showed that all the proteins, except Cef3, Fau1, Tsa1b, Mis1 and Erg10, take part in metabolic, cellular and localization processes **(Figure 5D)**, that only Atp2, Rps6, Rpl8 and Rps4 are involved on the constitution of a protein-containing complex and a cellular anatomical entity **(Figure 5E)**, and that all the identified proteins are mainly involved on catalytic and binding activities **(Figure 5F)**.

Finally, the analysis of the interaction between the identified proteins showed that they are involved in the GTP hydrolysis and joining of the 60S ribosomal subunit, the formation of a pool of free 40S subunits, and the glyoxylate and dicarboxylate metabolism. It should be highlighted that the Tsa1b, detected in gene and protein expression studies, is not included in any of the mentioned categories **(Figure 6)**.

**Figure 6.**
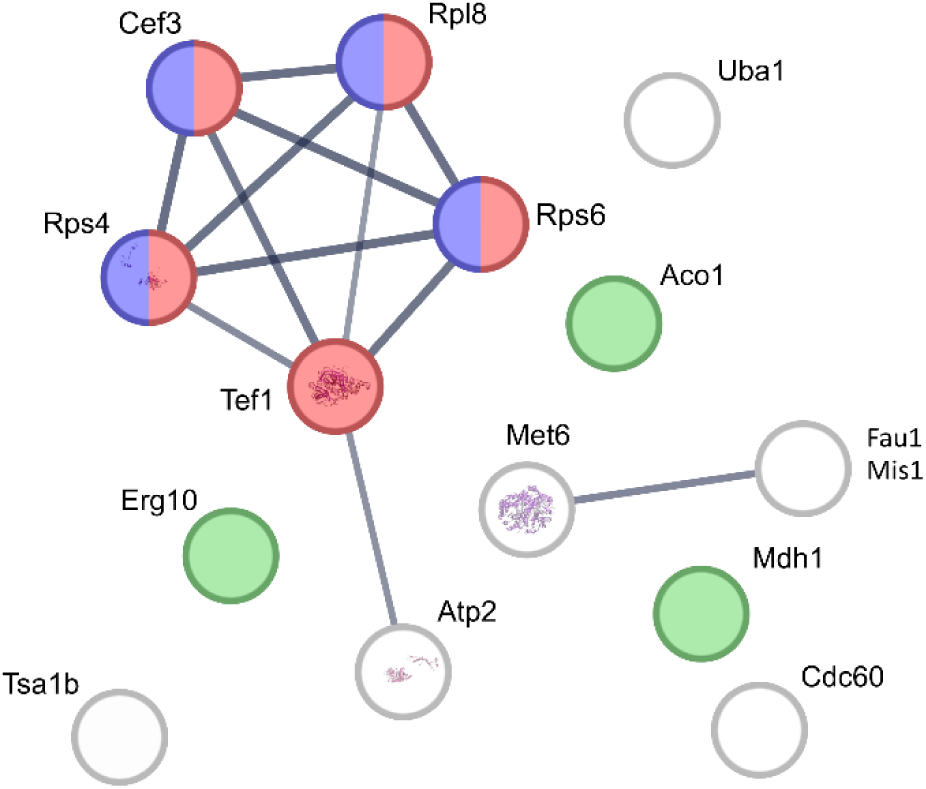
Diagram of the interaction between the specifically expressed and overexpressed proteins detected when *Candida auris* was grown on conditions of oxidative stress. The links between the proteins are indicated. Proteins involved on GTP hydrolysis and joining of the 60S ribosomal subunit, formation of a pool of free 40S subunits, and glyoxylate and dicarboxylate metabolism are indicated by red, purple, and green spots, respectively. Proteins not involved in any of these categories are not colored. STRING bioinformatic tool was used for the analysis of the interaction of the identified overexpressed proteins.

### Study of the antigenicity of the *C. auris* Tsa1b peroxiredoxin

The antigenicity of the Tsa1b peroxiredoxin was first studied by two-dimensional western blot (2D-WB) using a pool of sera obtained from patients with disseminated *C. auris* infection **(Figure 7A)**. Due to the high and diffuse reactivity observed in the acidic and high Mr area, probably caused by glycoproteins, the membranes were treated with sodium metaperiodate to oxidize and remove carbohydrates from glycoproteins, and the 2D-WB was repeated **(Figure 7B)**. In this way, four spots (4, 7, 9 and 13) previously identified as overexpressed under oxidative stress were also detected as antigens, corresponding to Tsa1b, Met6 and Mdh1. Moreover, as the reactivity observed for Tsa1b was weak, its antigenic role was individually verified by electroelution (P1) and WB was repeated against the same pool of sera, using another non-reactive electroeluted spot (P2) as control **(Figure 7C-D)**.

**Figure 7.**
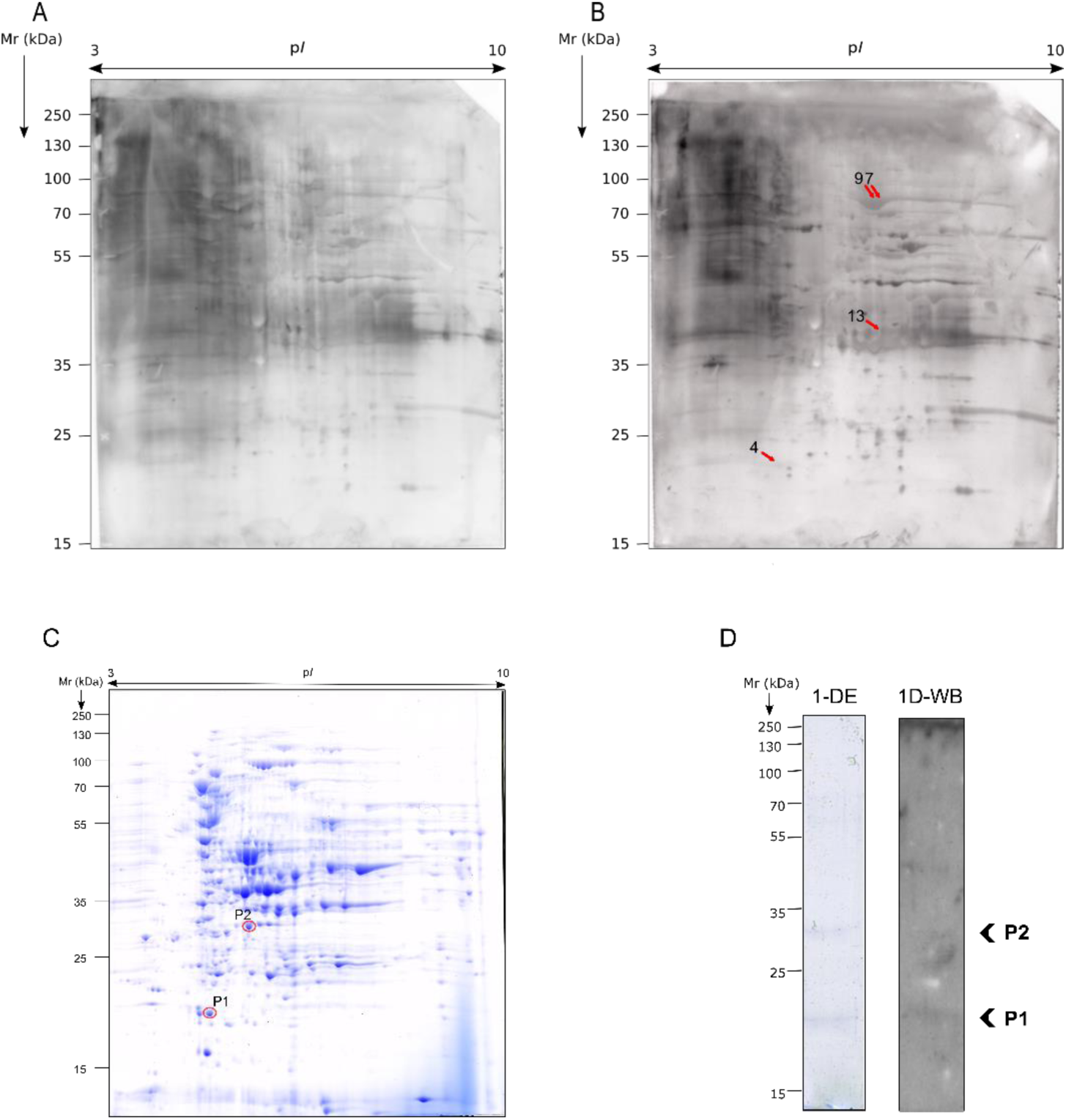
Detection of antigens recognised by human sera and present on *Candida auris* proteome obtained under conditions of oxidative stress and analysis of Tsa1b protein immunogenicity. 2D-WB images before (A) and after oxidation (B) of glycoproteins. *C. auris* CECT13225 was grown in the presence of 8 mM of H2O2. Dots marked with red arrows correspond to overexpressed immunogenic proteins; point 4 corresponds to thioredoxin-dependent peroxiredoxin (Tsa1b), points 7 and 9 to 5-methyltetrahydropteroyltriglutamate-homocysteine S-methyltransferase (Met6) and point 13 to malate dehydrogenase (Mdh1). Location of electroeluted spots on a 2-DE gel where P2 was used as a negative control and the protein in spot P1 corresponds to Tsa1b peroxiredoxin (C), and 1-DE and 1D-WB images of the electroeluted spots using pooled sera from infected patients (D) are shown.

### Relevance of *C. auris* Tsa1b peroxiredoxin on the growth capacity under environmental stresses and the cell wall composition

Given that the analysis of the gene and protein expression showed a higher expression of the Tsa1b in the presence of 8 mM of H_2_O_2_, a *C. auris* knock-out strain was obtained by CRISPR-Cas9 to further understand its role on fungal growth and virulence. For that, the *C. auris* DSM 105992 strain, previously used by other authors (Ennis et al., 2021; Rybak et al., 2020), had to be used because no deletion mutants were obtained with the *C. auris* CECT 13225 isolate used until now. In consequence, first it was corroborated that *TSA1B* was also overexpressed in this new strain in response to the oxidative stress by RT-qPCR in presence of 15 mM of H_2_O_2_, as this strain showed to be more resistant than CECT 13225 to this stress **(Figure S1A-B)**.

Then, *C. auris ΔTSA1B* mutant and complemented *ΔTSA1B::TSA1B* strains were generated, and the growth capacity of these strains under the presence of different stressors was studied **(Figure 8A)**. The results showed that the *C. auris ΔTSA1B* strain was more sensitive than the WT and complemented strains to the oxidative stress.

**Figure 8.**
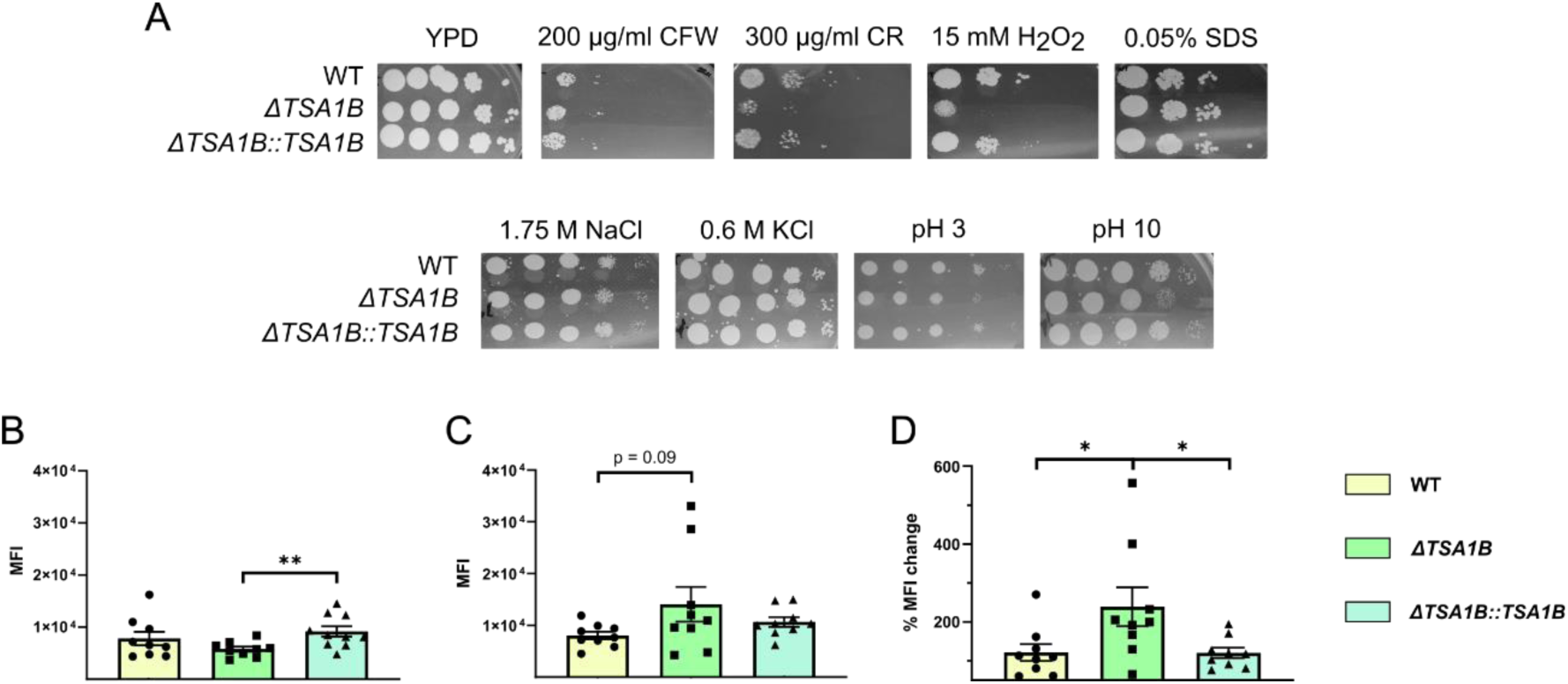
Characterization of the *Candida auris ΔTSA1B* strain generated by CRISPR-Cas9. Growth of *Candida auris* wild type (WT), *ΔTSA1B* and *ΔTSA1B::TSA1B* strains when grown on Yeast Peptone Dextrose (YPD) plates supplemented with stressor compounds (A). Ten µl of 1:10 serial dilutions starting at a density of 5 x 10^7^ *C. auris* yeasts/mL were plated on YPD plates supplemented with 200 µg/mL CFW, 300 µg/mL CR, 0.05% SDS, 15 mM H2O2, 1.75 M NaCl, 0.6 M KCl and with pH 3 or pH 10. Images were taken after incubation at 37°C for 24-48 hours. The most representative replicate of each condition is shown. Mean Fluorescence Intesity values recorded for β-glucan binding 100 mg/mL of aniline blue of *C. auris* WT, *ΔTSA1B* and *ΔTSA1B::TSA1B* strains in absence (B) and presence (C) of 15 mM of H2O2, and change percentage of the MFI values (D). Data was analysed by unpaired t-test. * p < 0.05, ** p < 0.01. Mean and standard error of mean values are indicated.

It was highlighted that concerning cell wall stressors, *C. auris ΔTSA1B* showed higher sensitivity to the β-glucan binding CR, but not to the quitin binding CFW. To verify these results, a semi-quantification of cell wall components chitin and β-glucan was performed in absence and presence of oxidative stress by fluorescence measurement using flow cytometry. In concordance to the result observed in YPD plates, no differences were observed in the detected chitin amounts of the *C. auris ΔTSA1B* strain in absence and presence of oxidative stress when compared with the WT and *ΔTSA1B::TSA1B* strains (data not shown). On the contrary, the quantification of β-glucan showed that the *C. auris ΔTSA1B* strain had the lowest signal conferred by β-glucans in absence of oxidative stress **(Figure 8B)**, and the highest in presence of 15 mM H_2_O_2_ **(Figure 8C)**. In order to measure the effect of oxidative estress in β-glucan detection, the increase in MFI values between absence and presence of H_2_0_2_ conditions was calculated. This revealed a statistically significant increase of β-glucan presence in the *C. auris* deletion mutant strain when compared to the WT and complemented strains **(Figure 8D)**.

As the *C. auris ΔTSA1B* seemed to have a modified cell wall composition in the presence of oxidative stress, it could have also an altered fungal resistance to antifungal drugs. Therefore, AFST was performed with fluconazole, amphotericin B and micafungin. However, no differences were recorded for any compound between *C. auris* WT and *ΔTSA1B* strains **(Table 2)**.

**Table 2.** Minimal Inhibitory Concentration (MIC) values (mg/l) of *C. auris* WT, *ΔTSA1B* and *ΔTSA1B::TSA1B* strains to fluconazole, amphotericin B and micafungin. On the case of fluconazole and micafungin, MIC50 values are indicated and for amphotericin B MIC90 values are shown. Results were obtained after incubation at 37°C for 24 hours.

| Strain | Concentration (mg/l) |  |  |
| --- | --- | --- | --- |
|  | MIC<sub>50</sub> Fluconazole | MIC<sub>90</sub> Amphotericin B | MIC<sub>50</sub> Micafungin |
| WT | 1-2 | 0,25-0,5 | 0,5 |
| ΔTSA1B | 1-2 | 0,25-0,5 | 0,5 |
| ΔTSA1B::TSA1B | 1-2 | 0,25-0,5 | 0,5 |

### Relevance of *C. auris* Tsa1b peroxiredoxin on the interaction with dendritic cells and macrophages

To investigate the role of the Tsa1b peroxiredoxin on the infectious process developed by *C. auris*, the host-fungus interaction was analysed. For that, the murine DC line DC1940, as well as BMDMs were infected with *C. auris* WT, *ΔTSA1B* or *ΔTSA1B::TSA1B* strains to quantify viable yeasts, ROS levels produced by immune cells, cytotoxicity levels on cells, and the uptake capacity. Furthermore, the genes of *C. auris* that were shown to be overexpressed in the presence of oxidative stress in this study were further analyzed when immune cells were infected.

When DCs and BMDMs were infected, a lower number of CFUs were recovered from knock-out *C. auris* compared to WT and complemented strains, the differences being statistically significant with respect to WT **(Figures 9A and 9G)**. Furthermore, on both immune cell types infected with *C. auris ΔTSA1B* strain, a higher level of ROS was detected than in those infected with WT and complemented strains **(Figures 9B and 9H)**, being the differences statistically significant on DCs. Finally, no differences were found in the cytotoxicity levels induced **(Figures 9C and 9I)**.

**Figure 9.**
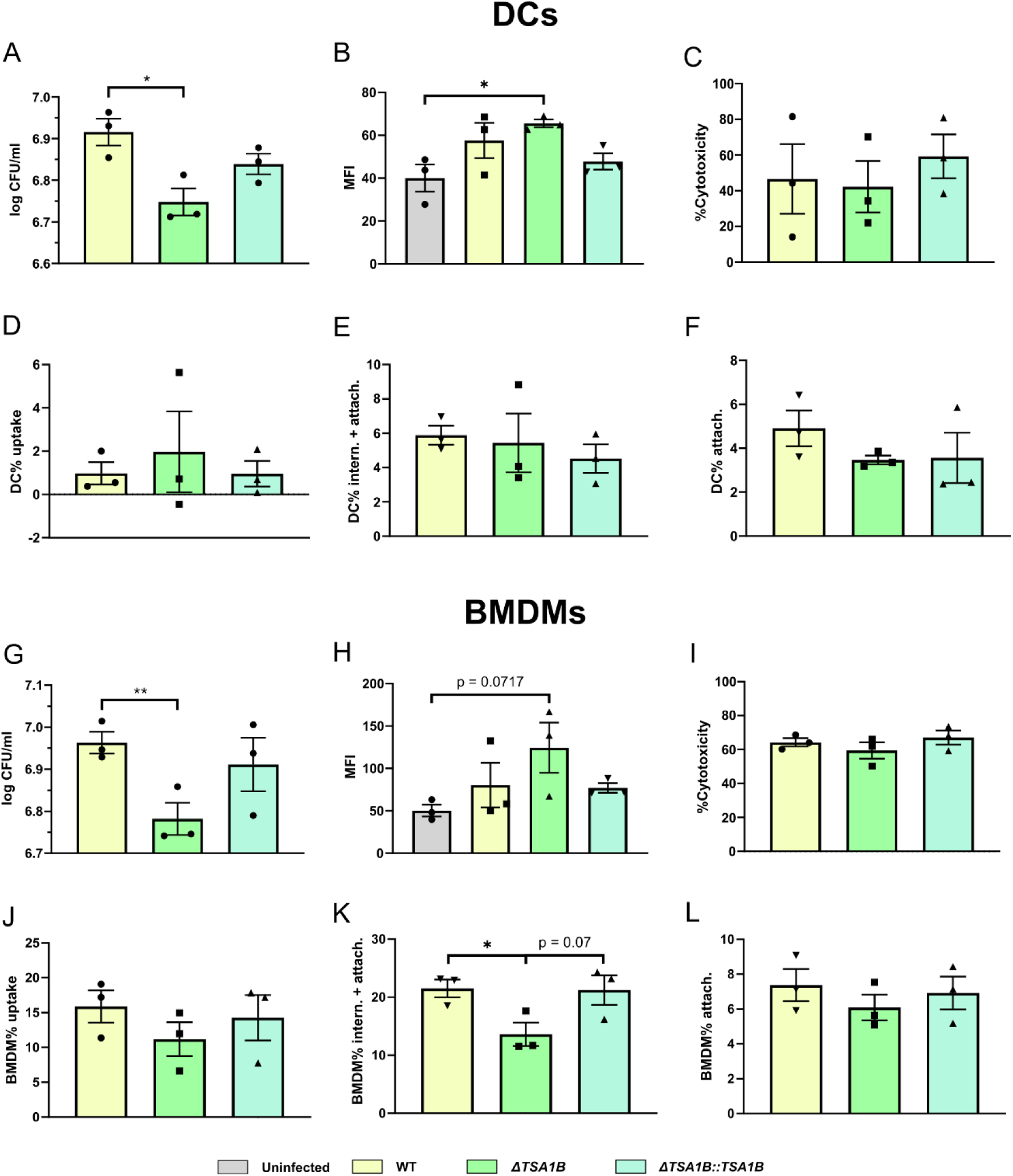
Response of murine DC1940 cells and BMDMs to *Candida auris* infection. Following a 4-hour co-incubation of DCs or BMDMs and *C. auris* yeast cells at a MOI of 5, viable *C. auris* cells recovered from DCs (A) and BMDMs (G) and mean fluorescence intensity (MFI) values detected by ROS production of DC cells (B) and BMDMs (H) are presented. The fraction of dead DCs (C) and BMDMs (I) because of *C. auris* infection was calculated after 24 hours. Finally, *C. auris* uptake capacity of DC1940 cells (D) and BMDMs (J), and internalization (intern.) and attachment (attach.) (E and K), and only attachment (F and L) of DC1940 cells and BMDMs proportion in contact with *C. auris* were measured after 30 minutes. Data was analysed by unpaired t-test. * p < 0.05. Mean and standard error of mean values are indicated.

After gating the collected events of DCs **(Figure S2)** and BMDMs **(Figure S3)** co-incubated with each *C. auris* strain, the uptake capacity was measured. While no differences were reported for DCs **(Figure 9D)**, a lower uptake when BMDMs were infected with the mutant strain than with the WT and complemented strains was observed, although the differences were not statistically significant **(Figure 9J)**. Specifically, in DCs, no differences were found, neither in the measure of the internalization and attachment (37°C condition) **(Figure 9E)** nor only attachment (4°C condition) **(Figure 9F)**, suggesting that *C. auris* Tsa1b peroxiredoxin is not involved in these processes. However, the separate analysis of the internalization and attachment **(Figure 9K)**, or only the attachment **(Figure 9L)** of *C. auris* cells to BMDMs, showed a significantly reduced amount of BMDMs interacting with *C. auris ΔTSA1B* strain on the 37°C condition, which might indicate a reduced internalization and attachment of the *C. auris ΔTSA1B* strain.

Then, the influence of the lack of Tsa1b on the expression of the rest of the previously detected overexpressed genes (*CAT1*, *SOD1*, *SOD6* and *CCP1*) was studied by RT-qPCR after infecting DCs or BMDMs for 2 and 4 hours. Using DCs, the results showed an overexpression of *CAT1*, *SOD1*, and *CCP1* genes on cells infected with the mutant strain at 2 hours **(Figure 10A)**. With BMDMs **(Figure 10C)**, underexpression of *SOD1* and not significant overexpression of *SOD6* were detected on *C. auris ΔTSA1B* strain at 2 hours of co-incubation. At 4 hours of co-incubation no significant differences were observed in any of the two models used **(Figures 10B and 10D)**.

**Figure 10.**
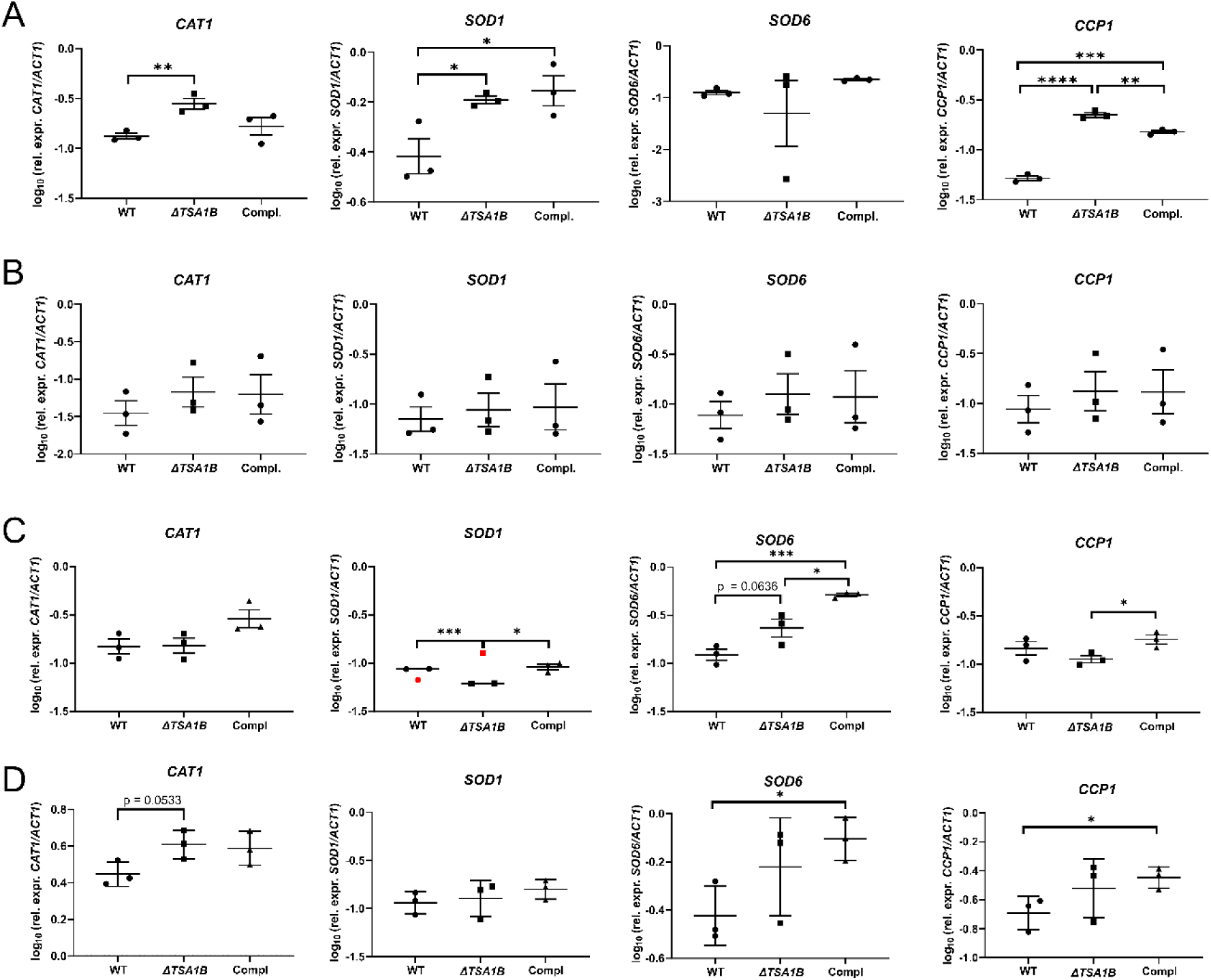
Gene expression analysis of *Candida auris* genes related to oxidative stress response on DC and BMDM infection models. After infection of DCs (A, B) and BMDM (C, D) cells with *C. auris* WT, *ΔTSA1B* and *ΔTSA1B::TSA1B* strains, RT-qPCR of *CAT1*, *SOD1*, *SOD6* and *CCP1* genes was performed. Gene expression analysis was carried out after co-incubation times of 2 hours (A, C) and 4 hours (B, D) of *C. auris* and host immune cells. Data was analysed by unpaired t-test. * p < 0.05, ** p < 0.01, *** p< 0.001, **** p < 0.0001. Mean and standard error of mean values are indicated. Outliers were detected with GraphPad outlier calculator, marked in red and removed from the statistical analysis.

### Study of the implication of Tsa1b on *C. auris* virulence *in vivo*

To determine the consequences of the lack of this protein on the virulence of the yeast, *G. mellonella* larvae and mice were infected. At seven days post-infection, 6.25% of *G. mellonella* larvae infected with the *C. auris ΔTSA1B* strain survived, whereas none of the larvae infected with the parental and complemented strains reached this time point. Furthermore, a delayed development of the infectious process was observed, recording greater survival percentages on the first two days of infection **(Figure 11)**.

**Figure 11.**
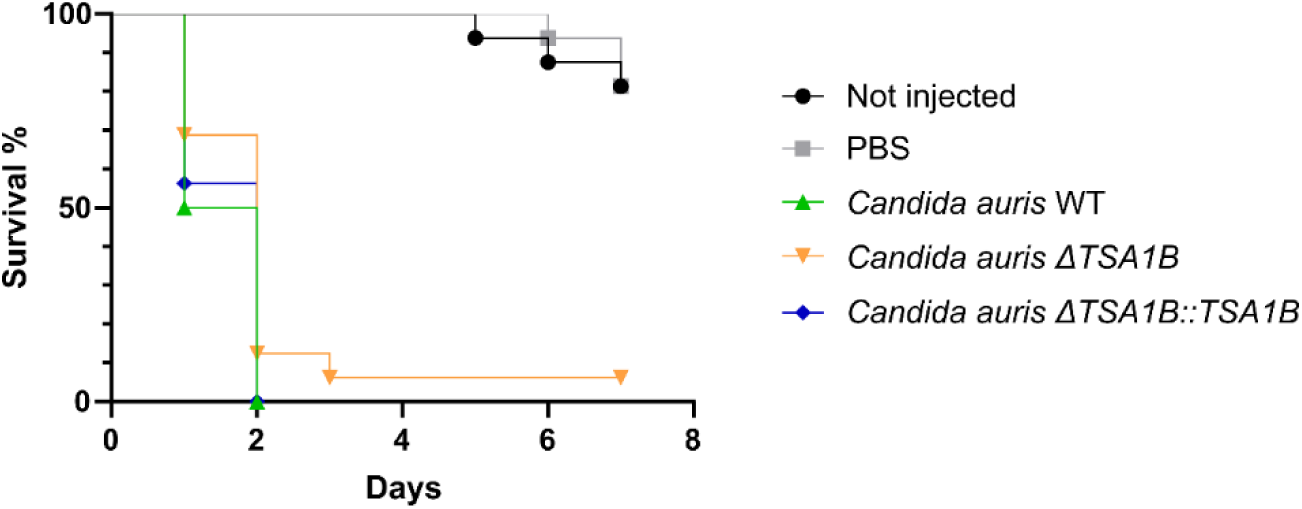
Survival curve of *Galleria mellonella* larvae infected with 2.5 x 10^6^ cells of *Candida auris* WT, *C. auris ΔTSA1B* and *C. auris ΔTSA1B::TSA1*B strains. Survival of *G. mellonella* was measured during 7 days. The most representative replicate is shown. Data was analysed by log-rank Mantel-Cox test for survival analysis.

To corroborate these results, a murine model of disseminated candidiasis was also used. The determination of the fungal load showed that the kidneys and brain were the most affected organs **(Figure 12A)**, the mutant strain showing lower fungal loads than WT strain in kidneys (p = 0.0677) and than complemented strain in brains (p = 0.0572). In addition, a greater spleen weight was detected on mice infected with *C. auris* WT and *ΔTSA1B::TSA1B* strains than with *ΔTSA1B* strain, being differences statistically significant between WT and *ΔTSA1B* strains. Finally, mice infected with WT and *ΔTSA1B* strains and sacrificed earlier than 28 days post-infection had lower spleen weight than those of the same sex that reached the end of the experiment **(Figure 12B)**.

**Figure 12.**
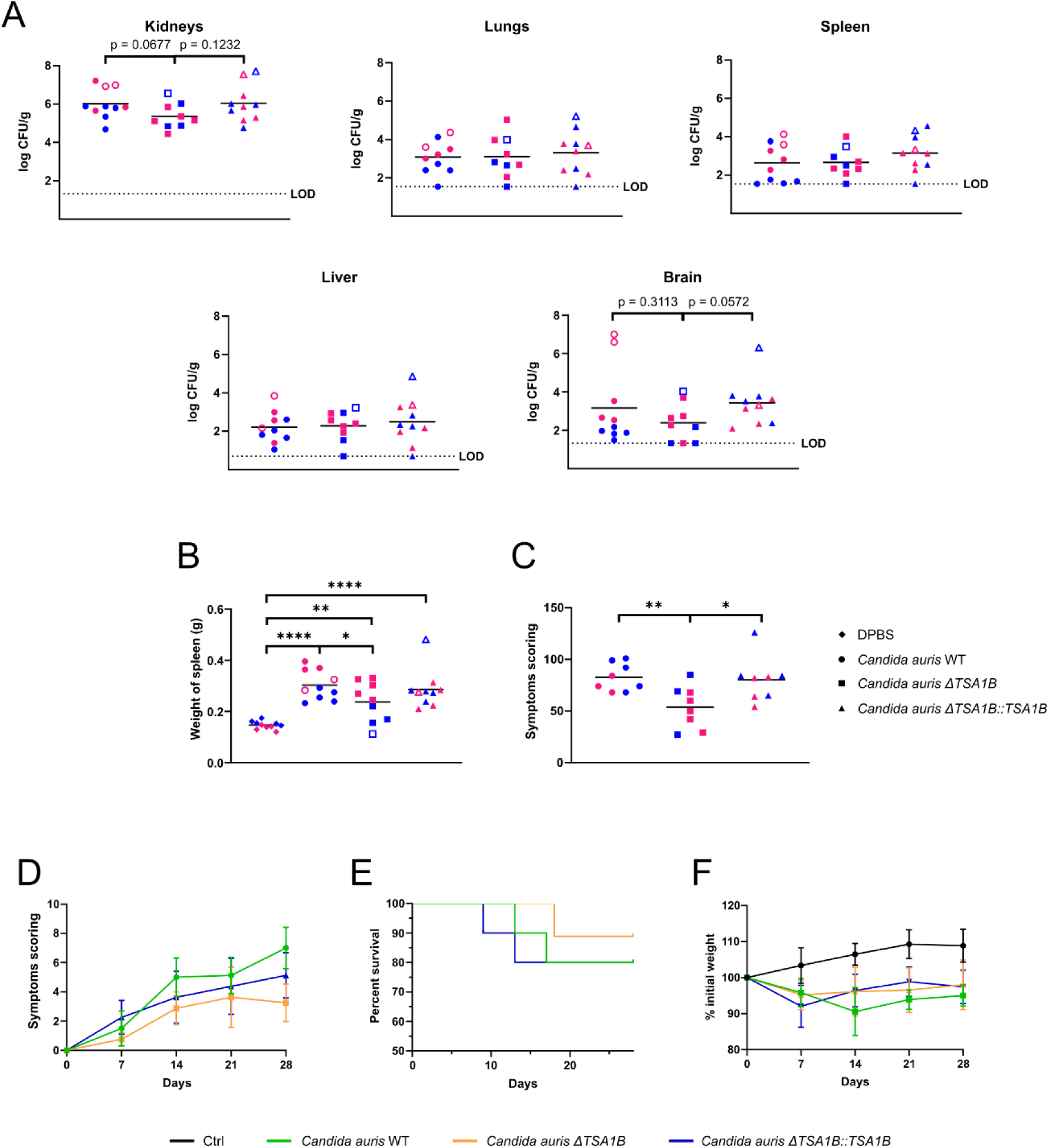
Study of the implication of *Candida auris* Tsa1b peroxiredoxin on the pathogenicity and infectious process developed by the yeast. Immunocompetent mice were intravenously injected with DPBS or 5 x 10^7^ yeasts/animal of *C. auris* WT, *C. auris ΔTSA1B* and *C. auris ΔTSA1B::TSA1B* strains. Fungal load of kidneys, lungs, spleen, liver and brain of the infected mice (A), weight of the spleens (B) and the cumulative value of the recorded symptoms (C) are shown. Pink and blue symbols correspond to female and male mice, respectively. Filled symbols refer to the mice reaching the end of the experiment, and empty symbols to those sacrificed at earlier time-points due to reaching humane end-points. Evolution of the presented symptoms (D), survival curve (E) and evolution of the weight (F) of the infected mice are also shown. The group infected with *C. auris ΔTSA1B* strain had an n = 9, as one mouse died because of the anaesthetics. The limit of detection (LOD) for each organ is indicated. All data below the LOD, including 0 values, were censored at this limit. Data was analysed by unpaired t-test. * p < 0.05, ** p < 0.01, *** p < 0.001, **** p < 0.0001.

Regarding the signs indicative of infection **(Table S4),** the mice infected with *C. auris ΔTSA1B* strain showed less cumulative symptoms than those infected with *C. auris* WT and *ΔTSA1B::TSA1B* strains **(Figure 12C)**. On the other hand, a slower, but not significant, appearance of the symptoms was noted on the mice infected with *C. auris ΔTSA1B* strain **(Figure 12D)**.

Additionally, greater survival rate was documented for the mice infected with the deletion strain than for those infected with the WT or complemented strains, although not significant differences were observed. In fact, among the mice sacrificed before the 28 days post-infection, those infected with the WT or complemented strains reached out the humane endpoint earlier than mice infected with the *ΔTSA1B* strain **(Figure 12E)**. Finally, mice infected with the *C. auris ΔTSA1B* strain showed a slower weight change in the first 14 days post-infection than those infected with the WT and complemented *C. auris* strains, but differences were not significant **(Figure 12F)**. It should be noted that the weight gain observed at later times corresponded to the sacrifice of mice that reached the humane endpoint, which lost the most weight.

The comparative study between sexes showed that females were more susceptible than male to *C. auris* systemic infection. Specifically, among the mice infected with the *C. auris* WT strain significant differences on kidneys and brain were found. However, it is important to consider the impact on the results of the high CFU/g values associated with the female animals euthanized at early time points on brain tissues **(Figure S4A)**. Regarding the weight of the spleens, those of females infected with either *C. auris* WT or *C. auris ΔTSA1B* strains suffered a greater weight gain than spleens of male **(Figure S4B)**, which seems to indicate a stronger immune response. Concerning the symptoms and weight loss no differences were detected (data not shown).

The microscopic examination of the kidneys and brain did not determine whether the *C. auris ΔTSA1B* strain exhibited different virulence than the *C. auris* WT or *ΔTSA1B::TSA1B* strains. Overall, a considerable number of fungal cells were detected in the tubular epithelium of *Candida*-exposed kidney samples **(Figures 13B–C)**, whereas no positive reaction was observed in the tubules of control mice **(Figure 13A)**. In consequence, an immune reaction was observed in all *Candida*-exposed groups, but the immune cell granulomatous inflammation was highly variable in terms of both number and size, with no evidence of a clear pattern regarding the infectious fungal strain or mouse sex. Of note, characteristic granulomatous inflammation was clearly observed with important granulomatous infiltrations **(Figures 13F-H)** on the kidneys of mice infected with either of the three *C. auris* strains. However, only in the kidneys of mice infected with the WT strain severe structural alterations were observed in both tubules and glomeruli **(Figure 13F and inset)**.

**Figure 13.**
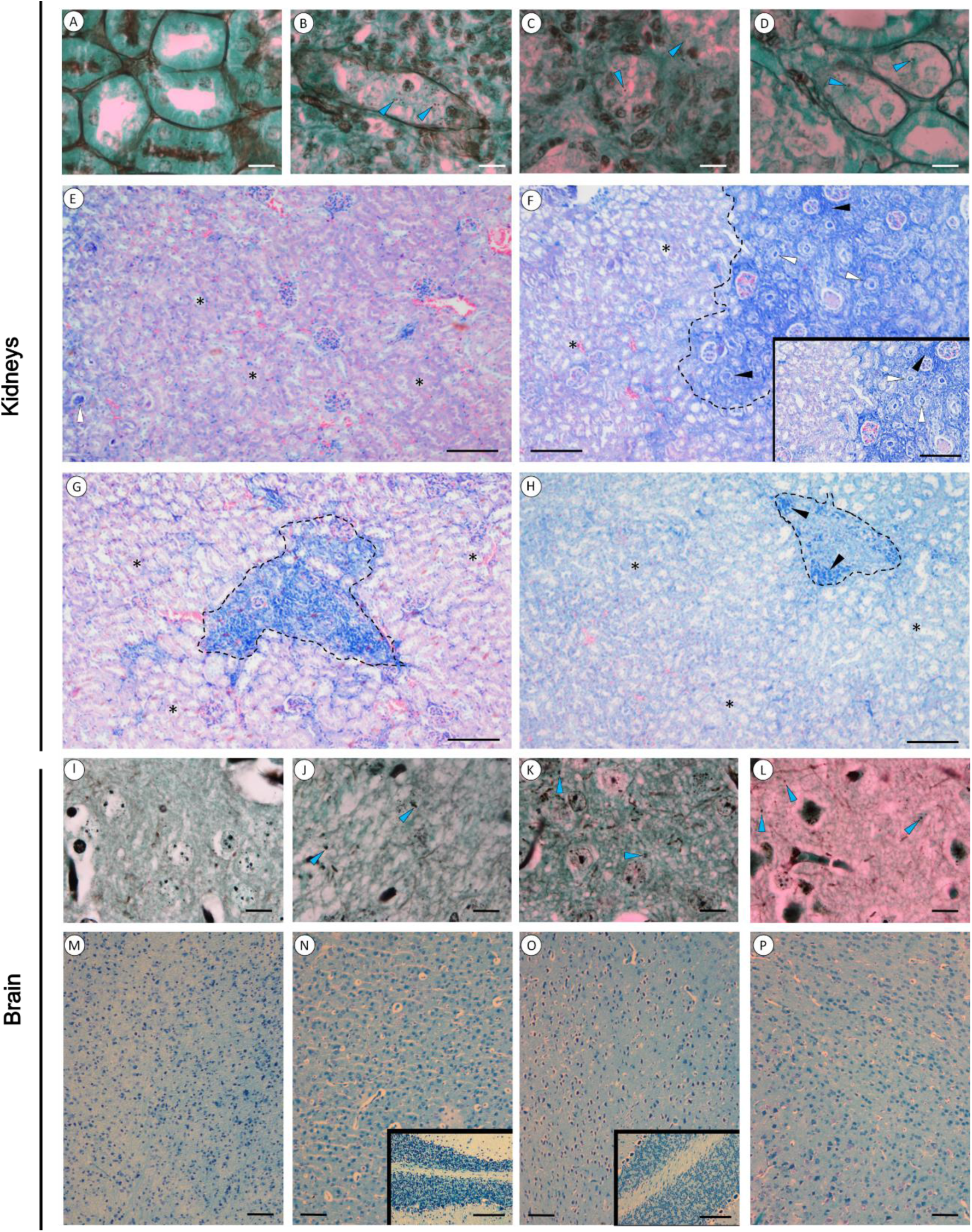
Kidney (A-H) and brains (I-P) of control (A, E, I and M), wild type (B, F, J and N), mutant (C, G, K and O) and complemented (D, H, L and P) *Candida auris* strains exposed mouse stained with Gomori Methenamine Silver (A-D and I-L) and Giemsa (E-H and M-P) stains. In the GMS stain of kidneys (A-D) and brain (I-L) some fungi can be observed in the exposed animals (blue triangles). Note the immune reaction occurred in the kidneys of the three infected groups (F-H, limited by the black dashed line) and the presence of immune cells (F-H, black triangles). In the Giemsa stained sections, the released material in the kidney tubules of exposed mice (F and inset, white triangles) is evident in comparison to the normal shaped tubules (F-H, asteriks). Note the similar cellular structure and pattern of the brain tissue in all samples (M-P) and the normal structure of cerebellum (insets of N and O) in *Candida* exposed mice. Scale bar = 10 µm (A-D and I-L) and 100 µm (E-H and M-P).

In the brain tissues of the analysed animals, only a few fungal particles were visualised in the brain tissues of infected animals with GMS staining **(Figures 13J–L)**, while no *C. auris* cells were observed in the uninfected mice **(Figure 13I)**. Furthermore, in the brain tissues recovered from control mice and those exposed to *Candida* did not reveal any differences in tissue structure or cell pattern **(Figures 13M–P)**. In addition, no structural alterations were detected in cerebellum tissue collected from *Candida*-exposed animals, as shown in the insets of **Figure 13**.

## DISCUSSION

*Candida auris* has several distinct characteristics that set it apart from other pathogenic *Candida* species, which include non-aggregative and aggregative growth phenotypes (Borman et al., 2016), as well as resistance to various environmental stresses, including oxidative stress (Day et al., 2018; Heaney et al., 2020). This is particularly significant because many disinfectants are based on peroxide solutions (Spivak and Hanson, 2018) and ROS play a role in host immune cell-mediated pathogen killing (Bassoy et al., 2021; Herb and Schramm, 2021; Martinvalet and Walch, 2022; Yang et al., 2013). Therefore, the aim of this study was to enhance the understanding of the factors involved in the infectious process caused by *C. auris*.

Concerning the growth capacity of *C. auris* clinical isolates under different stress conditions, greater resistance than *C. albicans* and *C. haemulonii* was observed, as previously reported by Day *et al*. (2018) and Heaney *et al*. (2020). More specifically, *C. auris* was less susceptible than *C. albicans* and *C. haemulonii* to the stresses caused by the chitin-binding CFW, the β-glucan-binding CR, the oxidative stress produced by H_2_O_2_, and the SDS-induced cell membrane damage.

Then, the response of *C. auris* in gene and protein level to oxidative stress was investigated, since ROS molecules produced by various immune cells can damage lipids, proteins, and DNA of fungal cells, and, consequently, *Candida* species need to return the redox balance to homeostasis (Dantas et al., 2015; Pradhan et al., 2017). Specifically, overexpression of the genes *CAT1*, *TSA1B*, *CCP1*, *SOD1*, and *SOD6* was found after 8 hours of oxidative stress incubation, *CAT1*, *TSA1B*, *CCP1* and *SOD1* being also overexpressed after 16 hours. Remarkably, two of the five genes that were previously overexpressed, *CAT1* and *SOD6*, showed underexpression at 24 hours, which might indicate a potential restoration of the cells to homeostasis. However, at 24 hours of oxidative stress, *HYR1*, *SSK1*, and *SOD2* were also found to be underexpressed. It is hypothesized that these genes are overexpressed during incubation periods shorter than 8 hours, as reported by some studies for *HYR1* and *SSK1* genes (Chauhan et al., 2003; Manfredini et al., 2004; Miramón et al., 2014; Vázquez et al., 2017), and, later, to restore the basal state of *C. auris*, their expression levels are reduced.

In addition, the study of the fungal proteome allowed the identification of fifteen proteins overexpressed under oxidative stress (Uba1, Cef3, Cdc60, Fau1, Tsa1b, Mis1, Met6, Aco1, Atp2, Erg10, Mdh1, Rps6, Rpl8, Rps4, and Tef1). Given that oxidative stress is known to cause protein damage (Martinvalet and Walch, 2022), the presence of enzymes involved in protein synthesis seems reasonable. In fact, Cef3, Rps4, Rps6, Rpl8, Tef1 and Uba 1 have been identified as necessary for replacing damaged proteins and proteins involved in ribosome structure (Dever et al., 2018; McGrath et al., 1991; Mirzaei and Regnier, 2006). Similarly, proteins implicated in the methionine biosynthesis (Met6, Fau1, and Mis1) are also relevant, since many proteins suffer methionine substitutions in their aminoacid sequences to protect cell and proteins from ROS (Campbell et al., 2016; Luo and Levine, 2009). Some of the proteins were involved in the glyoxylate cycle (Aco1, Mdh1, and Erg10), and as it happens in *Paraccocidioides lutzii* under oxidative stress (de Arruda Grossklaus et al., 2013), they could be overexpressed to keep the levels balanced, since they might suffer degradation (Dirmeier et al., 2002). Moreover, the overexpression of the enzyme Erg10 might indicate higher production of ergosterol, which is also common in other yeasts under oxidative stress conditions (Higgins et al., 2003; Lin et al., 2021; Montañés et al., 2011). The homology of the identified proteins with other fungal species and human proteins showed that the highest percentage of similarity was found with *C. haemulonii*, which is the closest phylogenetic fungus within the genus *Candida* (Satoh et al., 2009). However, due to significant differences with proteins found in *C. albicans*, *A. fumigatus*, and *H. sapiens*, some of these proteins may be of interest for potential therapeutic and diagnostic applications.

Based on both genetic expression and proteomic analyses conducted under oxidative stress, Tsa1b peroxiredoxin was the only molecule identified in both studies. Regarding the biological function of this enzyme, the *Saccharomyces cerevisiae* cytosolic yeast isoform is essential for resistance to exogenous H_2_O_2_ (Garrido and Grant, 2002). Furthermore, it triggers the recruitment of chaperones during protein aggregate formation caused by ROS molecules, which facilitates the transport of these aggregates to autophagosomes (Hanzén et al., 2016). Additionally, it has been demonstrated that Tsa1b provides protection against prion formation and structural alterations that might happen during oxidative stress (Doronina et al., 2015).

Given its relevance, Tsa1b antigenicity was confirmed using a pool of sera obtained from patients with disseminated candidiasis caused by *C. auris*, and a deletion mutant strain for this protein was constructed by CRISPR-Cas9 to further investigate its function during the interaction with the host. The generation of the mutant in the *C. auris* CECT 13225 isolate was not possible, probably because this strain belongs to clade III (Ruiz-Gaitán et al., 2018), which, along with clade IV isolates, presents a lower homologous recombination efficiency (Mayr et al., 2020). Therefore, *TSA1B* overexpression in the oxidative stress condition was verified in the clade I *C. auris* DSM 105992 strain, and deletion and complemented strains were constructed using this strain.

The *ΔTSA1B* strain exhibited a higher susceptibility to the cell wall stressor agent CR and oxidative stress generated by H_2_O_2_ compared to the WT and complemented strains. Similarly, Urban *et al*. noted a comparable pattern upon deletion of the *C. albicans* Tsa1b protein (2005). The semi-quantification of the cell wall component β-glucan showed that the higher susceptibility of the mutant *ΔTSA1B* strain to cell wall stressor CR could be linked with its high oxidative potential (Liu et al., 2021) and the considerable higher detection of β-glucan observed after exposure to oxidative stress.

Due to the relevance of *C. auris* Tsa1b peroxiredoxin on the response developed by the yeast against the oxidative stress response, and on the cell wall composition of the fungus, its implication was studied on the interaction with immune cells, such as DCs and BMDMs. In consequence, the *C. auris ΔTSA1B* strain was shown to be more susceptible to killing by DCs and BMDMs than the WT and *ΔTSA1B::TSA1B* strains, which might be a consequence of the higher ROS amount found in cells infected with *ΔTSA1B* than in those infected with the WT strain. This finding might be due to increased ROS generation by cells infected with the mutant or decreased ROS degradation by this strain, which could be related to the absence of Tsa1b peroxiredoxin (Wood et al., 2003). The cytotoxicity levels of the fungus on DCs and BMDMs were unaffected by the absence of the Tsa1b.

Regarding the uptake capacity, lower amounts of BMDMs, but not of DCs, that had uptaken *C. auris ΔTSA1B* compared to WT and *ΔTSA1B::TSA1B* were detected. In this context, BMDMs are known to express the β-glucan-binding dectin-1 receptor at higher levels than DCs (Municio et al., 2013). Therefore, this differential uptake capacity of BMDMs may be related to changes in cell wall composition due to the absence of Tsa1b protein, as it has been previously reported the presence of this protein in the cell wall of *C. albicans* hyphal cells (Urban et al., 2005). Linked to this, Tsa1b protein could be implicated in the the cell wall composition of *C. auris,* which would be supported by the lesser amount of β-glucans detected on *ΔTSA1B* strain in absence of oxidative stress. Hence, more research is needed to corroborate whether the *C. auris ΔTSA1B* strain has an altered cell wall composition and how this can modulate the host immune response, since β-glucan increase has been linked with important virulence traits of *C. albicans* (Bekirian et al., 2024). Additionally, overexpression of other genes related to the response to oxidative stress such as *CAT1*, *SOD1*, *SOD6*, and *CCP1* was detected in the *C. auris ΔTSA1B* strain after 2 hours of co-incubation with host immune cells, which may be linked to an attempt to compensate the absence of the Tsa1b under the oxidative stress generated by the cells, as observed with the upregulation of antioxidant enzymes in the oxidative stress response of *C. albicans ΔTSA1B* strain (Urban et al., 2005).

The study of the role of this protein during the infectious process *in vivo* showed that, the survival rates of animals infected with the *C. auris ΔTSA1B* strain were higher than with the WT strain, but without significant differences, as previously described with the same mutant in *C. albicans* (Urban et al., 2005). In fact, the progression of the infectious process in mice infected with the *ΔTSA1B* strain appears to be slower and less virulent, given the decreased weight loss and fungal loads, decreased spleen size, and fewer symptoms exhibited. As previously stated in this study, the absence of Tsa1b peroxiredoxin may negatively influence the capacity of the fungus to neutralise ROS generated by host immune cells, which may account for the slower rate of spread of the infection and increased killing efficiency of *C. auris*.

Concerning to differences between sexes, our study revealed that female mice were more affected by *C. auris*, even though females can develop stronger immunological responses (Lu et al., 2021). In fact, compared to male mice, female mice infected with *C. auris* WT showed a higher fungal load in the kidneys and brain, and a larger spleen. However, these differences between sexes might be due to the lower body weight of female than male mice and the use of the same infective dose for both sexes.

The histological examination of kidney and brain tissues from mice infected with the WT, *ΔTSA1B,* and *ΔTSA1B::TSA1B* strains did not show any significant difference. The similar damage caused by the three strains used may result from the compensatory overexpression of *CAT1*, *SOD1*, *SOD6*, and *CCP1* genes as well as other relevant genes involved in ROS detoxification. Therefore, further experiments with infection times shorter than 28 days could be interesting to perform to determine the essential role of the *C. auris* Tsa1b in infection progression.

In conclusion, this study has provided new insights into the response developed by *C. auris* to oxidative stress. In this context, relevant genes and proteins involved in the defence against ROS have been identified, with particular interest in Tsa1b peroxiredoxin since it has been detected as overexpressed in both gene and protein expression analyses. The generated *C. auris ΔTSA1B* strain exhibited a greater susceptibility to oxidative and cell wall stresses, as well as to DCs and BMDMs, a significant β-glucan amount increase in presence of oxidative stress, and a slower progression of the infectious process in *in vivo* animal models. However, further studies are needed to determine the potential use of this protein as a target for novel therapeutic or diagnostic approaches.

### Data Availability Statement

The raw MS data associated with this manuscript was submitted to the ProteomeXChange database and is available under the identifier PXD051894.

## Author Contributions

M.A., O.R-E., L.A.-F., L.A.-D., L.M.-S. and U.P.-C. carried out the *in vitro* experiments, and M.A., O.R-E., I.B., A.A. and A.Ra.-Ga. performed the experiments with mice. M.A. and B.Z. accomplished the histological study. A.Ru.-Ga. and J.P. obtained the *C. auris* isolates and the sera from human patients. S.L.-L, A.R., A.A. and A.Ra.-Ga. conceived the experiments and supervised the work. M.A. and A.Ra.-Ga. wrote the manuscript. All authors have actively contributed in reviewing the manuscript and gave approval to the final version of it.

## ACKNOWLEDGMENTS

Animal experimentation and mass spectrometry and flow citometry analysis were performed in the services provided by the SGIKER facility at the UPV/EHU.

