## Supplementary files for "The oxidative stress response-related peroxiredoxin Tsa1b of *Candida auris* functions as a virulence factor that promotes infection"

*[1]Department of Immunology, Microbiology and Parasitology, Faculty of Science and Technology, University of the Basque Country (UPV/EHU), Leioa, Spain; [2]Department of Immunology, Microbiology and Parasitology, Faculty of Pharmacy, University of the Basque Country (UPV/EHU), Vitoria-Gasteiz, Spain; [3]CBET Research Group, Department of Zoology and Animal Cell Biology, Faculty of Science and Technology, Research Centre for Experimental Marine Biology and Biotechnology PIE, University of the Basque Country (UPV/EHU), Leioa, Spain; [4]Microbiology Department, University and Polytechnic La Fe Hospital, Valencia, Spain; [5]Section of Immunology, Vetsuisse Faculty, University of Zürich, 8057, Zürich, Switzerland; [6]Institute of Experimental Immunology, University of Zürich, 8057, Zürich, Switzerland*

Corresponding author: Aitziber Antoran.

Address: [1]Department of Immunology, Microbiology and Parasitology, Faculty of Science and Technology, University of the Basque Country (UPV/EHU), Leioa, Spain

Telephone number: +34 946 01 5407

### SUPPLEMENTARY MATERIAL

**Table S1. Oligonucleotides used throughout the study.** Primers used for the qPCR expression analysis of *C. auris* genes related to the oxidative stress response, crRNAs used for the transformation of *C. auris* strains (their sequence and function is included and targeted PAM regions are underlined), primers used for the construction of the cassettes for the generation of mutant and complemented *C. auris* strain. (70 bp microhomology regions are indicated with lower case letters) and primers used for the screening of the deletion mutant and complemented *C. auris* strain are shown.

**Gene expression analysis**

| Gene | Forward primer (5' → 3') | Reverse primer (5' → 3') | Used for analysis |
|---|---|---|---|
| *CAT1* | GTCATCTTGTTCTCCGACCGT | CCAGTTGCCGTCCTTTGTAGA | I, II |
| *SOD1* | TTGGCAGATCTGTGGTTGTCC | GACGCCAATAACACCACAAGC | I, II |
| *SOD2* | GGTGGTGCTTTGGATGTTGTC | CAAGTAGTAAGCGTGCTCCCA | I |
| *SOD6* | ATCTTCAACCCTTACCACGCC | GTCTGGATTTGACCGTGCTTG | I |
| *SOD6* | TGCCGAGTTCCCTATGTTTGG | TGACCTTGACTTTCTTGCCGT | II |
| *HOG1* | TGGTCTGTTGGCTGCATTTTG | GCCCAAGAGCTCTGTGATGAT | I |
| *MAD2* | CCGGTTGAGAGATGGGTGTTT | GCCTGAATCTCCCTCTGCATT | I |
| *TSA1B* | GTGTTGTTTGCCTCGACTGAC | GCAAGCAATGGGATGTTGACA | I, II |
| *GPX2* | TGCGTCTTTCTGCCAGCTTAA | TGGGTCTGCATTATCGCCATT | I |
| *SSK1* | CAAACGCTCACACTCGAATCC | CGCCCAAAGTGAAGTTCTTCG | I |
| *CCP1* | TACAGATCGGGCTATGACGGT | TTGCCTCTTACCATCCCACTG | I, II |
| *ACT1* | TACTCTGTGTGGATTGGTGGC | AACAATCGATGGACCGGACTC | I |
| *ACT1* | TGCTGTGTTCCCATCCATTGT | GATTGAGCCTCATCACCGACA | II |

**crRNA design**

| crRNA | Sequence (5' → 3') | Function |
|---|---|---|
| 1 | TGACGACGTGGGCTGCGAAACGG | Deletion mutant strain generation |
| 2 | GAAGTAGCATTGTAATTTTAAGG | |
| 3 | ACATAGACGCAGGGCCTTCGTGG | Complemented strain generation |
| 4 | GACTAAGGCATAGGTGTACTTGG | |

**Cassette construction**

| Primer | Primer name | Sequence (5' → 3') | Length (bp) | Function |
|---|---|---|---|---|
| 1 | Hyg Fw Cass | atgacgtcaaaaatactcgtgggttgctgatgtgaatgacctgcacatagacgcagggccttcgtggtgcGGTTTTCCCAGTCACGACGT | 90 | CaCO_HygR amplification for deletion mutant construction |
| 2 | Hyg Rv Cass | aggtgtacttggacttgaagaagacctgggactctacggcaaactggtagaagagcgtttccttggtcatGTGTGGAATTGTGAGCGGAT | 90 | |
| 3 | Compl TSA1B Fw Cass | atgcggttaattttaaaaaatctcccatgacgtcaaaaatactcgtgggttgctgatgtgaatgacctgcGACGCAGGGCCTTCGTCG | 88 | *TSA1B* gene amplification for complemented strain construction |
| 4 | Compl TSA1B Rv Hyg tail | taagttaaataaataattagataagggtggtaattattactatttacaatcaaaggtggtcctctgcaggATCTATAAAATTTATTTGTTGACCTTGCCG | 100 | |
| 5 | Compl Hyg Fw TSA1B tail | agaccatcaagcctgacgtgaagaactccaaggagtacttcggcaaggtcaacaaataaattttatagatCCTGCAGAGGACCACCTTTG | 90 | *hph* gene amplification for complemented strain construction |
| 6 | Compl Hyg Rv Cass | cgccaaatcgaagcacgctagttctgagaggcaccacaagaacgtggcccggcaccaagggtttgatgttCCATCATAAAATGTCGAGCGTCAAAAC | 97 | |

**Screening of deletion and complementated strains**

| Primer | Primer name | Sequence (5' → 3') | Length (bp) | Function |
|---|---|---|---|---|
| a | Scr TSA1B Fw | TGACCGCTATCATCCAGAAGC | 21 | *TSA1B* gene amplification |
| b | Scr TSA1B Rv | GCACACCTCACCGTACTTCT | 20 | |
| c | Scr Fw | GGGTTGCTGATGTGAATGACC | 21 | Screening of transformed colonies |
| d | Scr Rv | GTTCTGAGAGGCACCACAAGA | 21 | |

(I) Used in the gene expression study performed in absence and presence of oxidative stress

(II) Used in the gene expression study performed in the interaction with host immune cells

**Table S2. Staining panels used for DC1940, BMDM and *Candida auris* cells.** Antibodies used for the staining of different cells, the fluorescence channel on which they need to be visualized and the dilution factor applied are indicated.

| Cells | Antibody | Fluorescence channel | Dilution |
|---|---|---|---|
| **DC1940** | Anti-CD11c | PECy7 | 100 |
| | - | FITC | - |
| **BMDMs** | Anti-CD64 | PE | 100 |
| ***Candida auris*** | Chitin[a] | PB | 1000 |

**Table S3. Minimal Inhibitory Concentration (MIC) values (mg/l) of *Candida auris* isolates to fluconazole, amphotericin B and micafungin.** On the case of fluconazole and micafungin, $MIC_{50}$ values are indicated and for amphotericin B $MIC_{90}$ values are shown. Results were obtained after incubation at 37°C for 24 hours.

| | Concentration (mg/l) | | |
|---|---|---|---|
| **Isolate** | **$MIC_{50}$ Fluconazole** | **$MIC_{90}$ Amphotericin B** | **$MIC_{50}$ Micafungin** |
| **CJ-194** | >64 | 0,125 | 0,0625 |
| **CJ-195** | >64 | 0,125 | 0,03125 |
| **CJ-196** | >64 | 0,125 | 0,03125 |
| **CECT 13225** | >64 | 0,5 | 0,125 |
| **CECT 13226** | >64 | 0,125 | 0,0625 |

**Table S4. Scoring system used during the supervision of the mice in order to determine the relevant symptoms and their severity.** General and specific symptoms as well as the value (1-4) given to each of them are shown.

| General Symptom | Specific symptom | Value (1-4) |
|---|---|---|
| **Body weight** | Loss between 5-10% | 1 |
| | Loss between 10-20% | 2 |
| | Loss between 20-25% | 3 |
| | Loss equal to or greater than 25% | 4 |
| **Transient discomfort after injection** | | 1 |
| **Abnormal postures** | Hunched abdomen | 2 |
| | Stretching of the body | 2 |
| **Weakness or paralysis of the limbs** | | 4 |
| **Skin alterations** | Changes in skin consistency | 1 |
| | Ruffled hair | 2 |
| **Stool appearance** | Soft | 1 |
| | Diarrhoea | 2 |
| | Blood in stool | 3 |
| | Diarrhoea >48 hours | 3 |
| **Feeding and drinking** | Transient anorexia after injection | 1 |
| | Recurrent anorexia | 2 |
| | Not drinking | 3 |
| **Breathing** | Tachypnoea | 1 |
| | Dyspnoea | 2 |
| | Severe dyspnoea | 3 |
| **Neurological disturbances** | Head bobbing | 1 |
| | Leaning to one side | 1 |
| | Ataxia | 2 |
| | Jumping | 2 |
| | Complete loss of balance | 3 |
| **Behaviour** | Transient lethargy after injection | 1 |
| | Stereotypies | 1 |
| | Moderate change in behaviour and/or withdrawal from peers | 2 |
| | Persistent lethargy | 3 |
| | Reacts violently/vocalisation | 3 |
| **Physical parameters** | Distension of the abdomen | 2 |
| | Cachexia | 4 |
| | 20% increase in body circumference | 4 |

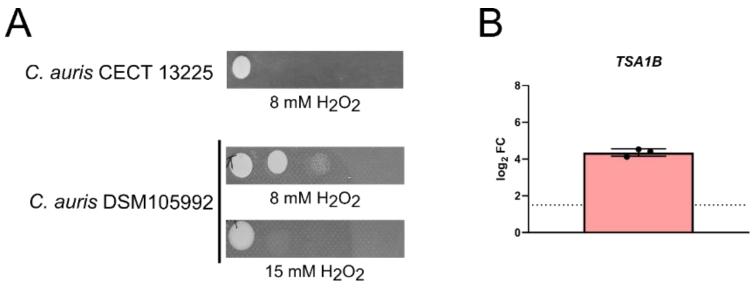


**Figure S1. Analysis of *Candida auris* DSM 105992 strain grown on conditions of oxidative stress.** Growth of *C. auris* CECT 13225 and DSM 105992 isolates on conditions of oxidative stress (A). To make sure the same degree of susceptibility as in CETC 13225 strain was achieved, 8 mM of $H_2O_2$ were used for CECT 13225 and 8 mM and 15 mM of $H_2O_2$ for the DSM 105992 isolate. Ten µl of 1:10 serial dilutions starting at a density of 5 x $10^7$ *Candida* yeasts/mL were plated on $H_2O_2$ supplemented SDA plates. Images were taken after incubation at 37 °C for 24 h. The most representative replicate of each condition is shown. qPCR analysis of DSM 105992 *TSA1B* gene after an incubation time of 8 hours on SDB supplemented with 15 mM of $H_2O_2$ (B). The fold change (FC) with respect to the control conditions and taking the reference expression of the constitutively expressed gene *ACT1* is represented. Threshold of 1.5 was chosen for the determination of the overexpression of the gene. Mean and standard error of mean values are indicated.

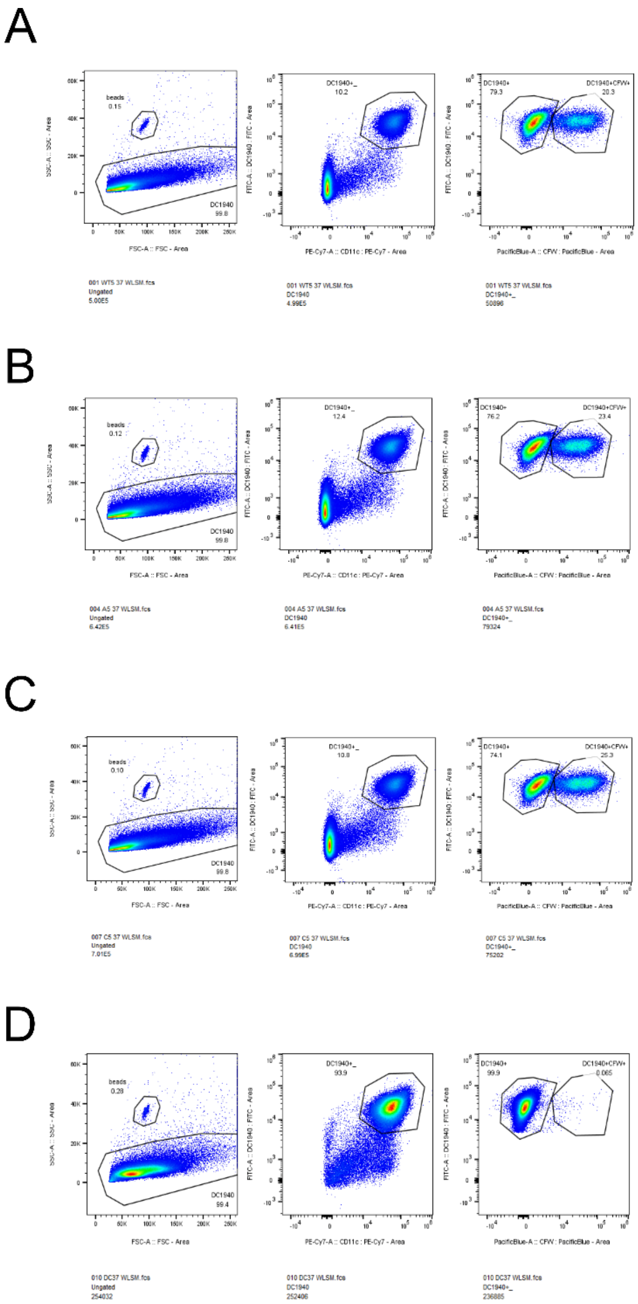


**Figure S2. Gating strategy followed for the determination of uptake capacity of DC1940 cells when incubated with *Candida auris*.** DC1940 cells co-incubated with *C. auris* WT (A), *ΔTSA1B* (B) and *ΔTSA1B::TSA1B* (C) strains, or without fungal cells (D). $GFP^{+}$ and $CD11c^{+}$ double positive events were considered DC cells. $CFW^{+}$ events were considered fungal cells. Analysis of the recorded events was made by FlowJo Software.

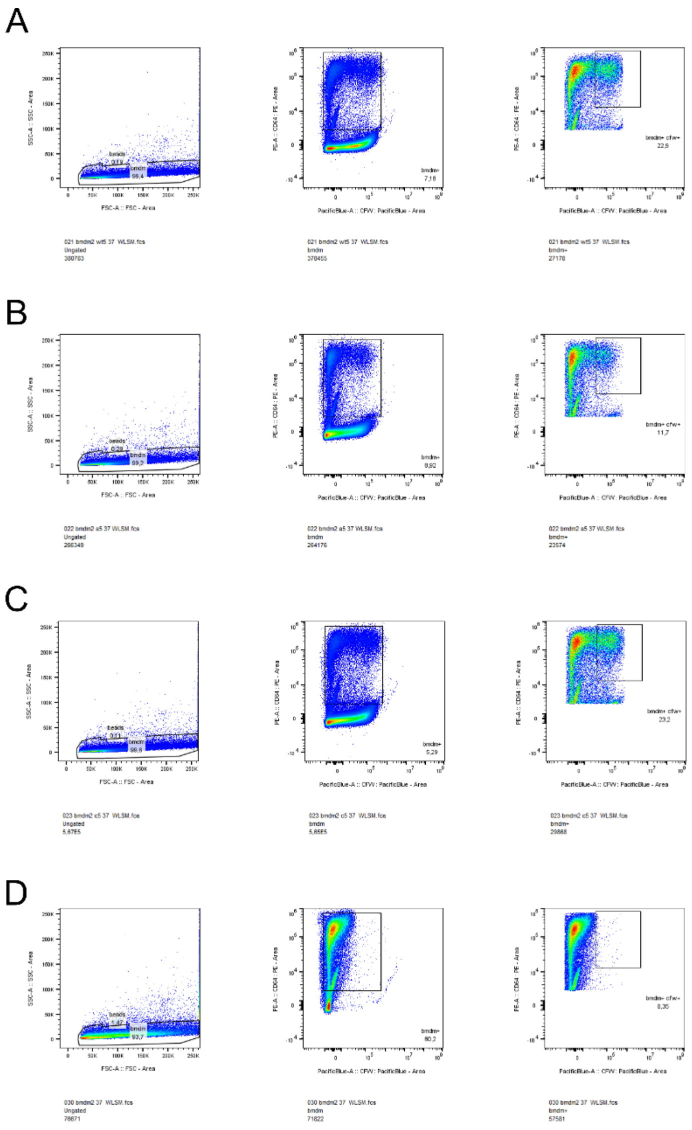


**Figure S3. Gating strategy followed for the determination of uptake capacity of BMDM cells when incubated with *Candida auris*.** BMDM cells co-incubated with *C. auris* WT (A), *ΔTSA1B* (B) and *ΔTSA1B::TSA1B* (C) strains, or without fungal cells (D). $CD64^{+}$ double positive events were considered BMDM cells. $CFW^{+}$ events were considered fungal cells. Analysis of the recorded events was made by FlowJo Software.

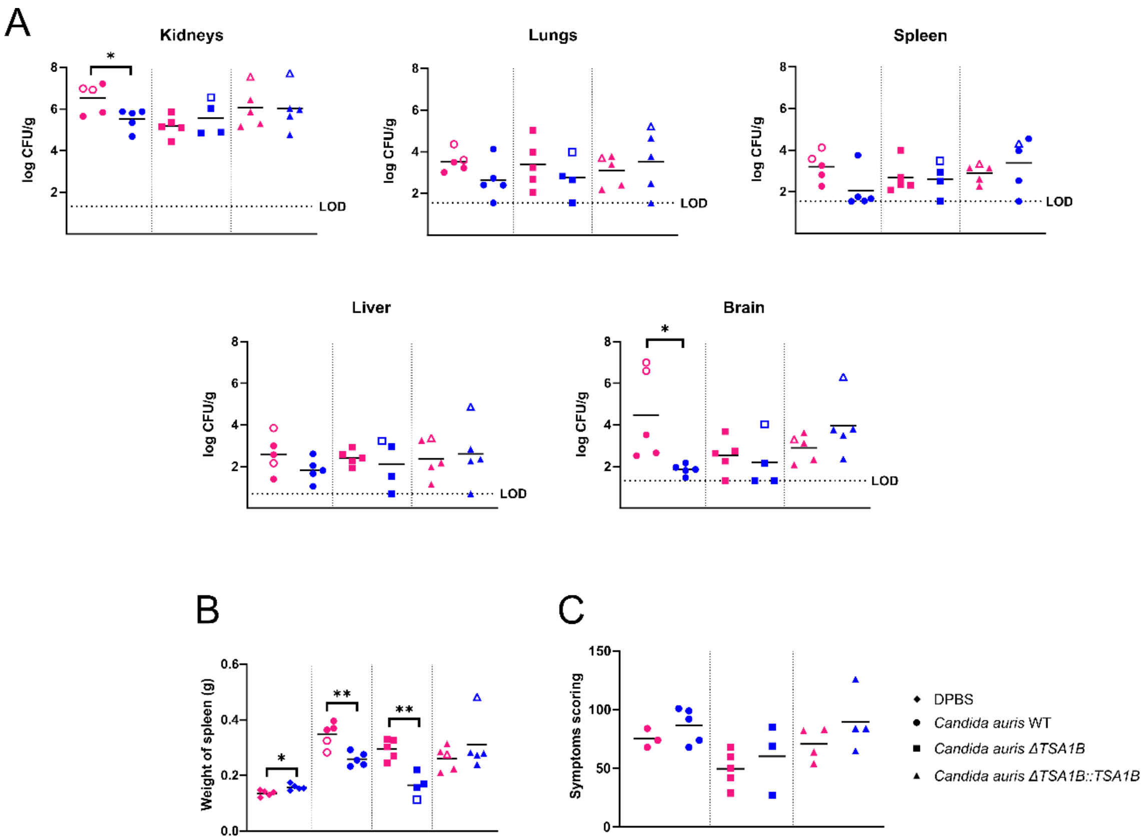


**Figure S4. Study of the implication of *Candida auris* Tsa1b peroxiredoxin on the pathogenicity and infectious process developed by the fungus on male and female mice.** Immunocompetent mice were intravenously injected with DPBS or 5 x $10^7$ yeasts/animal of *C. auris* WT, *C. auris ΔTSA1B* and *C. auris ΔTSA1B::TSA1B* strains. Fungal load of kidneys, lungs, spleen, liver and brain of the infected mice (A), weight of the spleens (B) and the cumulative value of the recorded symptoms (C) are shown. Pink and blue symbols correspond to female and male mice, respectively. Filled symbols refer to the mice reaching the end of the experiment, and empty symbols to those sacrificed at earlier time-points due to reaching humane end-points. The group infected with *C. auris ΔTSA1B* strain had an n = 9, as one mouse died because of the anaesthetics. The limit of detection (LOD) for each organ is indicated. All data below the LOD, including 0 values, were censored at this limit. Data was analysed by unpaired t-test. * $p < 0.05$, ** $p < 0.01$, *** $p < 0.001$, **** $p < 0.0001$.
